# Chronic pain exacerbates nicotine withdrawal severity in a sex-specific and dose-dependent manner

**DOI:** 10.64898/2026.04.16.719070

**Authors:** Brianna Graham, Tyler Nelson, Sheela Tavakoli, Laura O’Dell, Nii A. Addy, Deniz Bagdas

**Author notes:** **Corresponding Author:** Deniz Bagdas, DVM, PhD, Department of Psychiatry, Yale School of Medicine, 300 George Street, Suite 901, New Haven, CT, 06511, USA.

## Abstract

Chronic pain and nicotine use frequently co-occur, and individuals with chronic pain often experience greater difficulty quitting. Therefore, we examined nicotine withdrawal behaviors and analgesic-like effects in pain-naive and chronic pain conditions.

Adult male and female rats underwent chronic constriction injury or sham surgery. After pain establishment, rats received twice-daily subcutaneous nicotine (0.3 or 0.7 mg/kg) or saline for 14 days. 24 h after the final injection, withdrawal was assessed, including physical signs and anxiety-like behavior. Depressive-like responses were evaluated at 72 h. Pain sensitivity and nicotine’s analgesic-like effects were assessed throughout.

Chronic pain increased physical signs of withdrawal in both sexes, with greater effects in females. It also induced anxiety-like behavior in controls of both sexes. In rats with comorbid chronic pain and withdrawal, anxiety-like behavior was further enhanced in males, whereas females showed variable responses across assays, with increases or decreases depending on the test. Chronic pain induced depressive-like behavior in males but not in females. During withdrawal, depressive-like responses in males with chronic pain were not greater than those in the chronic pain alone group, while chronic nicotine exposure reduced depressive-like behavior in females. Nicotine produced acute analgesic-like effects that diminished over time in both pain-naive and chronic pain conditions, indicating tolerance. In pain-naive rats, repeated nicotine exposure induced mechanical hypersensitivity.

Chronic pain intensified nicotine withdrawal severity in a nicotine concentration- and sex-dependent manner. These findings highlight the importance of considering pain status and sex when developing effective cessation strategies, particularly for individuals with comorbid chronic pain.

**Summary:** Chronic pain exacerbates nicotine withdrawal severity. Chronic nicotine exposure induces pain hypersensitivity and tolerance to analgesic effects. These effects vary by nicotine concentration and sex.

## 1. Introduction

Chronic pain affects about 20 percent of U.S. adults [70]. It is accompanied by sensory, cognitive, and affective symptoms that contribute to both physical and emotional distress, including depression and anxiety [33,53,55]. Chronic pain is also often comorbid with other conditions. For example, individuals with chronic pain are significantly more likely to develop nicotine dependence than the general population [32,44,72]. Thus, nicotine use continues to be disproportionately high among individuals with chronic pain. Additionally, up to 60 percent of tobacco users also meet criteria for chronic pain [29], suggesting a clinically relevant comorbidity between chronic pain and nicotine use. Nicotine use remains one of the leading causes of preventable death [66,67], underscoring the need to understand the mechanisms driving this comorbidity.

The causal link between chronic pain and nicotine dependence remains poorly understood, particularly regarding the impact of pain on nicotine use. One possibility is that individuals use nicotine for its analgesic and mood-enhancing effects, driving dependence and complicating cessation efforts. While nicotine exhibits analgesic effects in humans [14,23] and antinociceptive properties in rodents [7,30], these effects diminish with prolonged exposure, indicating tolerance to its analgesic properties [35,41]. Moreover, nicotine withdrawal can heighten sensitivity to pain stimuli in both clinical and preclinical settings [6,11,42,48]. Withdrawal-related symptoms may therefore play a key role in maintaining nicotine use in individuals with chronic pain. Nicotine also has antidepressant-like properties [40,54,57], and many individuals report using nicotine to cope with pain-related distress [34]. Thus, the possibility exists that individuals with chronic pain become dependent on nicotine both to manage pain and to alleviate negative affective states. They may also experience more severe withdrawal symptoms during abstinence, which could facilitate a more rapid trajectory of nicotine dependence. Therefore, we aimed to examine the impact of chronic pain on the behavioral expression of the physical and negative affective states produced by nicotine withdrawal.

While both chronic pain and nicotine use are more prevalent and severe in women than in men [12,66,70], behavioral studies testing sex differences in their comorbidity remain limited, as these conditions have largely been studied separately. Women are more likely to experience chronic pain and show greater pain sensitivity than men [12,17,39,70]. While men tend to use e-cigarettes more for nicotine reinforcement, women are more likely to report use for mood regulation [51]. Although men smoke more cigarettes overall [43], women show greater sensitivity to nicotine concentration and dependence [45,47,49,50,58]. However, sex-specific behavioral studies on nicotine use in the context of chronic pain are lacking. To date, only one study has directly addressed this intersection, showing that chronic pain is more strongly associated with smoking in women than in men [1]. These findings underscore the need for research that captures sex-specific vulnerabilities in comorbid pain and nicotine use [8]. Thus, we examined how chronic pain influences nicotine withdrawal behaviors and evaluated nicotine’s effects on mechanical pain sensitivity in this comorbidity. Using a preclinical model, we assessed sex-specific behavioral outcomes following repeated nicotine exposure and spontaneous withdrawal in animals with established neuropathic pain.

## 2. Materials and methods

### 2.1. Animals

Adult female (240-255 g upon arrival) and male (300-330 g upon arrival) Sprague-Dawley rats (Charles River Laboratories, Wilmington, MA, USA) were used. Animals were housed in a temperature-controlled vivarium on a reverse 12-hour light/dark cycle (lights off at 7:00 AM) and acclimated for 4 days prior to experimental procedures. Initially, rats were group-housed (2 per cage) with *ad libitum* access to water and standard chow (#2018S, Envigo Teklad, Madison, WI). Following surgery, rats were housed individually. To maintain consistency with other operant-based studies in our lab, food restriction (6 g of pellets per 100 g body weight per day) was introduced on post-operative day 6, following an initial unrestricted feeding period to support recovery. All experiments were conducted during the light phase between 8:00 AM and 5:00 PM. All procedures were approved by the Yale University Institutional Animal Care and Use Committee and followed the National Institutes of Health Guidelines for the Care and Use of Laboratory Animals.

### 2.2. Chemicals

Nicotine tartrate salt (MP Biomedicals, Cat #153554) was dissolved in sterile 0.9% saline to prepare subcutaneous injection solutions at 0.3 and 0.7 mg/kg doses (calculated as free base). Solutions were filtered through sterile syringe filters, transferred to light-protected sterile glass vials, and stored at 4°C for up to 28 days. Injections were administered subcutaneously at a volume of 1 ml/kg body weight.

### 2.3. Surgical procedures

#### 2.3.1. Chronic constrictive injury-induced neuropathic pain model

To induce chronic neuropathic pain, we utilized the chronic constrictive injury (CCI) model of the sciatic nerve, based on the Bennett and Xie method with slight modifications [10,13]. Rats were initially divided into two groups: CCI and sham, with baseline pain sensitivity counterbalanced across groups. CCI surgeries were performed under 1-4% isoflurane anesthesia. Rats received preemptive analgesia with carprofen (5 mg/kg, subcutaneously) during surgery. The left sciatic nerve was exposed at the mid-thigh level using blunt dissection between muscle planes; no muscles were cut, and only fascia was separated. Two 5-0 silk sutures were loosely tied around the nerve approximately 1 mm apart. The incision site was then closed using 4-0 nylon sutures. Sham rats underwent identical dissection and closure procedures without nerve touch and ligation. Post-operative care included daily monitoring of body weight, hydration status, motor function, and general health. No analgesia was administered post-surgery, per approved IACUC protocol, to avoid interfering with pain assessments. Neuropathic pain was established within two weeks post-surgery, with pain localized to the ipsilateral hind paw.

### 2.4. Behavioral assessments

#### 2.4.1. Von Frey test

Mechanical paw withdrawal thresholds were assessed according to Dixon’s up-down method [24], with slight modifications [7]. Rats were placed in suspended wire-mesh chambers and allowed to habituate for 10 minutes. Calibrated von Frey filaments (Stoelting, Wood Dale, IL) were applied perpendicularly to the plantar surface of the hind paw. Testing was conducted within a force range of 1.479 to 125.892 g, which served as the lower and upper cutoff values. All testing was initiated with a 15.136 g filament for each rat and proceeded up or down based on the response. In the absence of a paw withdrawal response, a thicker filament corresponding to a stronger stimulus was applied. If a withdrawal response occurred, the next weaker filament was presented. On subsequent testing days, the initial filament was selected based on the threshold determined in the previous session. Baseline thresholds were assessed on both hind paws prior to surgery. To reduce potential sensitization during repeated testing, only the ipsilateral paw (relative to injury) was tested on nicotine testing days. The withdrawal threshold was defined as the lowest filament that elicited a consistent behavioral response, such as paw lift, licking, or shaking. A decreased paw withdrawal threshold on the ipsilateral side was interpreted as increased mechanical hypersensitivity consistent with mechanical allodynia associated with neuropathic pain. An increase in threshold following nicotine administration was interpreted as an anti-allodynic effect in rats with CCI surgery and anti-nociceptive effect in rats with sham surgery.

#### 2.4.2. Evaluation of somatic signs

To assess physical symptoms of spontaneous nicotine withdrawal, somatic signs were recorded during a 20-minute observation period. Rats were individually placed in empty polycarbonate cages (40.6 x 21.6 x 21.0 cm) and tested in pairs, with two cages positioned side by side on a table. A red lamp provided consistent lighting at approximately 30 lux, and a mirror was positioned behind the cages to enable full behavioral visibility. The following behaviors were scored: paw tremors, head shakes, body tremors, backing, scratching/itching, writhing, ptosis, curling, jumping, cheek tremors, teeth chattering, foot licking, floor licking, head/body swaying, and falling while standing. The total number of somatic signs was recorded for each rat, and data were reported as the average number of signs per group. A higher number of somatic signs was interpreted as increased physical withdrawal severity following nicotine exposure.

#### 2.4.3. Light/Dark transition test

To assess nicotine withdrawal-induced anxiety-like behavior, rats were tested in a custom-built light/dark box consisting of a clear, illuminated compartment and a connected black, enclosed compartment (30.5 x 30.5 x 30.5 cm each) with an arch-shaped opening between them. The light side was illuminated at 278 lux, and the dark side remained at 0 lux. Each rat was placed in the light compartment and allowed to explore freely for 10 minutes. Behavioral measures included time spent in the light compartment and the number of entries. A decrease in time spent in the light compartment was interpreted as indicator of anxiety-like behavior.

#### 2.4.4. Marble Burying Test

To assess nicotine withdrawal-induced anxiety-like behavior, rats were tested using the marble burying test. Each rat was placed in a novel cage containing 2 inches of bedding and 24 black marbles evenly spaced around the perimeter. The session lasted 30 minutes. At the end of the session, the number of marbles buried was recorded. A marble was considered buried if at least half of its surface was covered by bedding. Testing was conducted in a dark room with 0 lux illumination. An increased number of buried marbles was interpreted as an indicator of anxiety-like behavior.

#### 2.4.5. Forced swim test

To assess depressive-like behavior and stress reactivity, rats were placed in a clear polypropylene cylinder (30 cm in diameter, 60 cm high, with a water depth exceeding 40 cm) filled with water and a temperature maintained between 23 and 26 °C. The first session lasted 15 minutes and served as habituation or baseline exposure. The second session took place 24 hours later and lasted 6 minutes. Behavior during the second session was video recorded and later analyzed for immobility time. Increased immobility time was interpreted as a depressive-like behavioral response and heightened stress reactivity.

### 2.5. Experimental design

Rats underwent CCI or sham surgery on Day −14. Paw withdrawal thresholds were assessed weekly from Day −14 through Day 21 (Figure 1). Beginning on Day 0, animals received subcutaneous injections of saline, 0.3, or 0.7 mg/kg nicotine for 14 consecutive days (28 injections total: twice daily at 8:00 am and 4:30 pm for 13 days, with an additional single injection at 4:30 pm on Day 0 and at 8:00 am on Day 14). Von Frey testing was conducted before and after the injections on Days 0, 7, and 14 to evaluate acute antinociceptive effects of nicotine in sham rats and antiallodynic effects of nicotine in CCI rats. On Day 15 (24 h after the final injection), spontaneous withdrawal was assessed. Somatic signs were measured first, followed by the light dark transition test. Von Frey testing was performed next, and the marble burying test was conducted last. Forced swim test habituation was performed on Day 16, followed by forced swim test on Day 17. Animals were euthanized after von Frey testing on Day 21.

**Figure 1.**
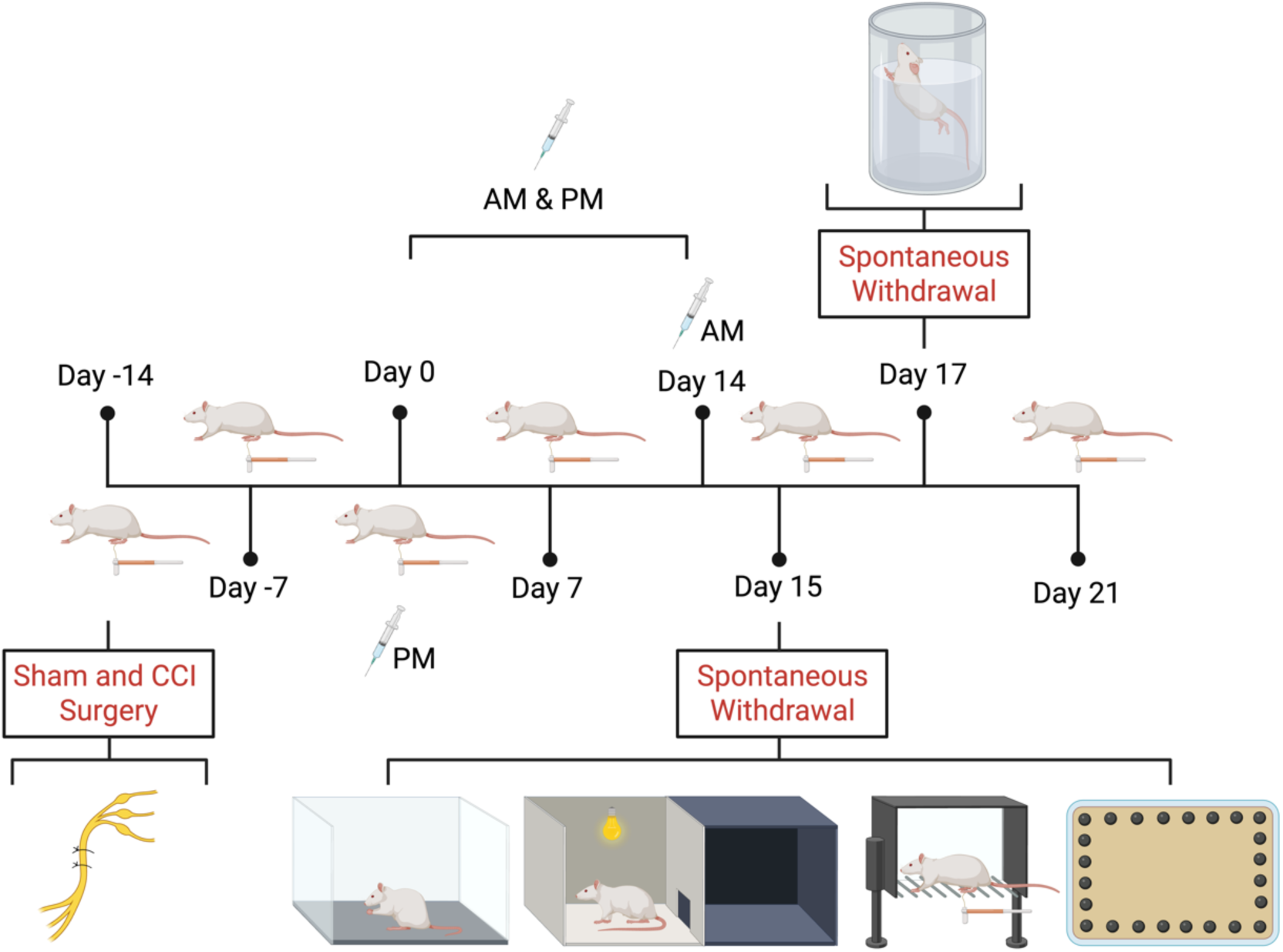
Experimental timeline of surgery, nicotine exposure, and behavioral withdrawal testing. Rats underwent chronic constrictive injury (CCI) of the sciatic nerve or sham surgery on Day −14. Pain sensitivity was assessed with von Frey test weekly from Day −14 to Day 21. Beginning on postoperative Day 0, animals received subcutaneous injections of saline, 0.3 mg/kg, or 0.7 mg/kg nicotine for 14 days (28 injections total; twice daily at 8:00 am and 4:30 pm, with a single injection at 4:30 pm on Day 0 and at 8:00 am on Day 14). Von Frey testing was conducted before and after injections on Days 0, 7, and 14 to assess antinociceptive effects in sham rats and antiallodynic effects in CCI rats. On Day 15 (24 h after the final injection), spontaneous withdrawal was assessed, including somatic signs and the light-dark transition test, followed by von Frey and marble burying tests. Forced swim test habituation and testing were conducted on Days 16 and 17, respectively. Animals were euthanized after von Frey testing on Day 21.

### 2.6. Blinding

All behavioral testing and video analyses were conducted by experimenters (BG, ST, TMN, DB) blind to treatment conditions; group coding was performed by an experimenter not involved in testing. A treatment key for groups was provided to DB for statistical analysis after behavioral scoring was completed.

### 2.7. Statistical analysis

Data were analyzed using GraphPad Prism 11 (GraphPad Software, San Diego, CA, USA). Prior to statistical testing, data were examined for normality and variance homogeneity. All datasets met the assumptions of normality and homogeneity of variance required for ANOVA. Two-way ANOVA was used for within-sex comparisons, with factors of surgery (CCI vs. sham) and nicotine dose. All data were also analyzed using three-way ANOVA to examine interactions among sex, surgery, and nicotine treatment. When significant main effects or interactions were found, post hoc comparisons were performed using Tukey’s multiple comparisons test. Significance was set at *p* < 0.05.

A two-way repeated measures (RM) analysis of variance with Greenhouse-Giesser correction was used to evaluate antinociceptive and antiallodynic effects of nicotine over the given time course. For RM analyses of paw withdrawal thresholds across time, data were analyzed in R (version 4.5.3; R Foundation for Statistical Computing, Vienna, Austria) using the afex package. A three-way RM ANOVA was conducted with chronic pain state and nicotine dose as between-subject factors and timepoint as a within-subject factor. To assess sex differences, an additional four factors repeated-measures ANOVA was performed including sex, chronic pain state, and nicotine dose as between-subject factors and timepoint as a within-subject factor. When the assumption of sphericity was violated, Greenhouse-Geisser corrections were applied. If Mauchly’s test indicated that the assumption of sphericity was met for all repeated-measures factors (all *p* > 0.45), Greenhouse-Geisser and Huynh-Feldt corrections were not applied.

One female rat was excluded due to post-surgical wound opening. One male rat was excluded due to >15% body weight loss in accordance with IACUC protocol. One male rat was excluded from the light-dark transition test due to video recording failure. In addition, for the light-dark transition test, rats that remained exclusively (>95%) on one side of the apparatus were excluded, and their data were removed from all behavioral analyses. To maintain a final sample size of 8 rats per group, a total of 7 additional rats (male and female) were tested across two separate cohorts. No male and female testing was conducted on the same day to control for potential confound. The final dataset included a total of 96 rats (3 nicotine dose groups by 2 surgical conditions by 2 sexes, with 8 rats per group).

## 3. Results

### 3.1. Impact of chronic pain on physical symptoms of nicotine withdrawal

On the withdrawal day, spontaneous withdrawal signs were quantified as first. Increased number of somatic signs were identified as physical symptoms of withdrawal. Nicotine induced physical symptoms of nicotine withdrawal in both male and female rats in a dose-dependent manner (Figure 2A and B). A two-way ANOVA revealed significant main effects of nicotine [F_(2, 42)_ = 69.06, *p* < 0.0001], chronic pain [F_(1, 42)_ = 27.10 *p* < 0.0001], and interactions between nicotine and chronic pain [F_(2, 42)_ = 3.941, *p* = 0.0270] for male rats (Figure 2A). In male sham rats, saline-treated ones showed minimal signs, while 0.3 and 0.7 mg/kg nicotine significantly increased withdrawal symptoms. In male CCI rats, similar effects were observed. While minimal signs were counted in saline-treated rats, nicotine at 0.3 and 0.7 induced significant physical symptoms of nicotine withdrawal. Post hoc analysis revealed that the same nicotine dose produced greater number of somatic signs in CCI rats compared to sham controls, indicating that chronic pain amplifies the physical symptoms of nicotine withdrawal in males.

**Figure 2.**
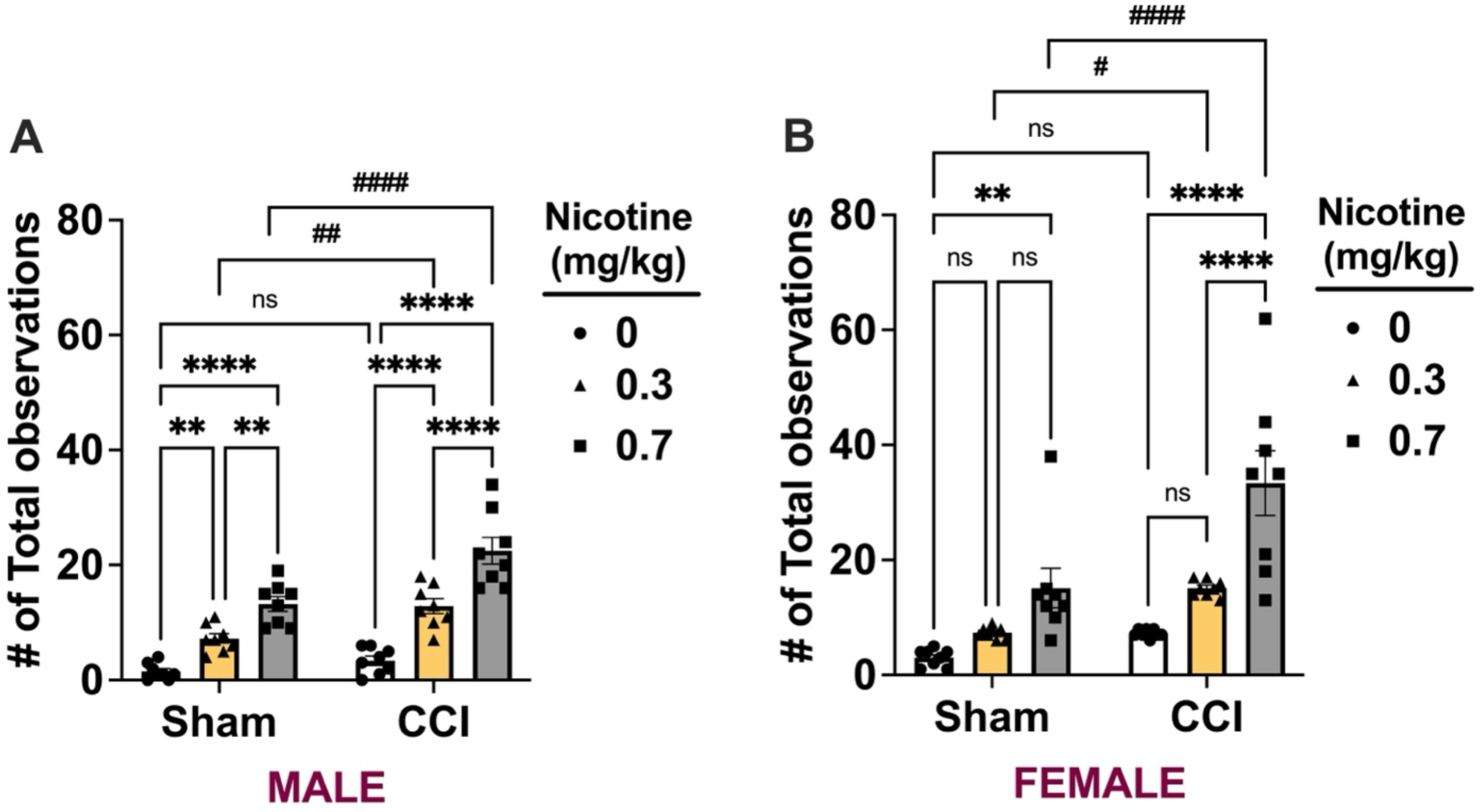
Impact of chronic pain on physical symptoms of nicotine withdrawal. Rats underwent sham or chronic constriction injury (CCI) surgery. Two weeks after surgery, animals received 14 days of subcutaneous injections of saline, 0.3 mg/kg, or 0.7 mg/kg nicotine. Approximately 24 h after the final injection, somatic withdrawal signs were assessed in male (A) and female (B) rats. Data represent the total number of somatic signs observed and are expressed as mean ± SEM. \**p* < 0.05 compared with saline; #*p* < 0.05 compared with sham.

In female rats, a two-way ANOVA revealed significant main effects of nicotine [F_(2, 42)_ = 25.94, *p* < 0.0001], chronic pain [F_(1, 42)_ = 20.74, *p* < 0.0001], and interactions between nicotine and chronic pain [F_(2, 42)_ = 3.609, *p* = 0.0358; Figure 2B]. While saline-treated female sham rats showed minimal signs, 0.7 mg/kg nicotine-treated female sham rats showed increased number of somatic signs (*p* < 0.01). In female CCI rats, saline induced several somatic signs, but it was not significantly different from sham controls (*p* > 0.05). Nicotine at 0.7 mg/kg dose significantly increased number of somatic signs compared to saline and 0.3 mg/kg nicotine. This suggests as evidence of physical symptoms of nicotine withdrawal. Post hoc analysis revealed that the same nicotine dose produced an even greater number of somatic signs compared to sham controls, indicating that chronic pain amplifies the physical symptoms of nicotine withdrawal.

Additional three-way ANOVA analyzed interactions between chronic pain state, nicotine dose, and sex effects (Supplementary Figure 1). We found significant main effects for nicotine dose [F_(2, 21)_ = 29.92, *p* < 0.0001], sex [F_(1, 21)_ = 8.202, *p* = 0.0093], and chronic pain [F_(1, 21)_ = 132.6, *p* < 0.0001] but no global interactions between these three [F_(2, 21)_ = 1.596, *p* = 0.2264]. There were significant interactions between nicotine dose and chronic pain state [F_(2, 21)_ = 21.28, *p* < 0.0001] but no interactions between nicotine dose and sex [F_(2, 21)_ = 0.669, *p* = 0.2125]. However, there was significant interactions between chronic pain and sex [F_(1, 21)_ = 6.376, *p* = 0.0197], which revealed sex-dependent effects in chronic pain conditions.

### 3.2. Impact of chronic pain on anxiety-like behaviors of nicotine withdrawal

Immediately after spontaneous withdrawal measurements, anxiety-like behaviors of withdrawal were evaluated in the light-dark transition test. Time spent in the light side of the box was used to index anxiety-like behavior; decreased time spent in the light side was identified as anxiety-like behavior of withdrawal. In males, nicotine decreased time spent in the light side in both sham and CCI in a dose dependent manner (Figure 3A). A two-way ANOVA revealed significant main effects of nicotine [F_(2, 42)_ = 15.99, *p* < 0.0001] and chronic pain [F_(1, 42)_ = 46.20, *p* < 0.0001], but no interactions between nicotine and chronic pain [F_(2, 42)_ = 0.4274, *p* = 0.6550] for male rats (Figure 3A). In male sham rats, while 0.3 mg/kg nicotine had no effect on time spent in light, 0.7 mg/kg nicotine significantly decreased time spent in light compared with saline-treated rats (*p* < 0.01), which indicates nicotine induced anxiety-like behavior. In male CCI rats, saline-treated rats showed a significant decrease in time spent in light compared with their sham controls, which suggests chronic pain induced anxiety-like behavior (*p* < 0.001). While 0.3 mg/kg nicotine had no effect on time spent in light, 0.7 mg/kg nicotine significantly decreased time spent in light compared with saline-treated CCI rats (*p* < 0.01), which indicates nicotine induced anxiety-like behavior. Planned post hoc analysis also revealed that the same nicotine dose produced greater anxiety-like behavior in CCI rats compared to sham controls, indicating that chronic pain amplifies the anxiety-like behavior of nicotine withdrawal in males.

**Figure 3.**
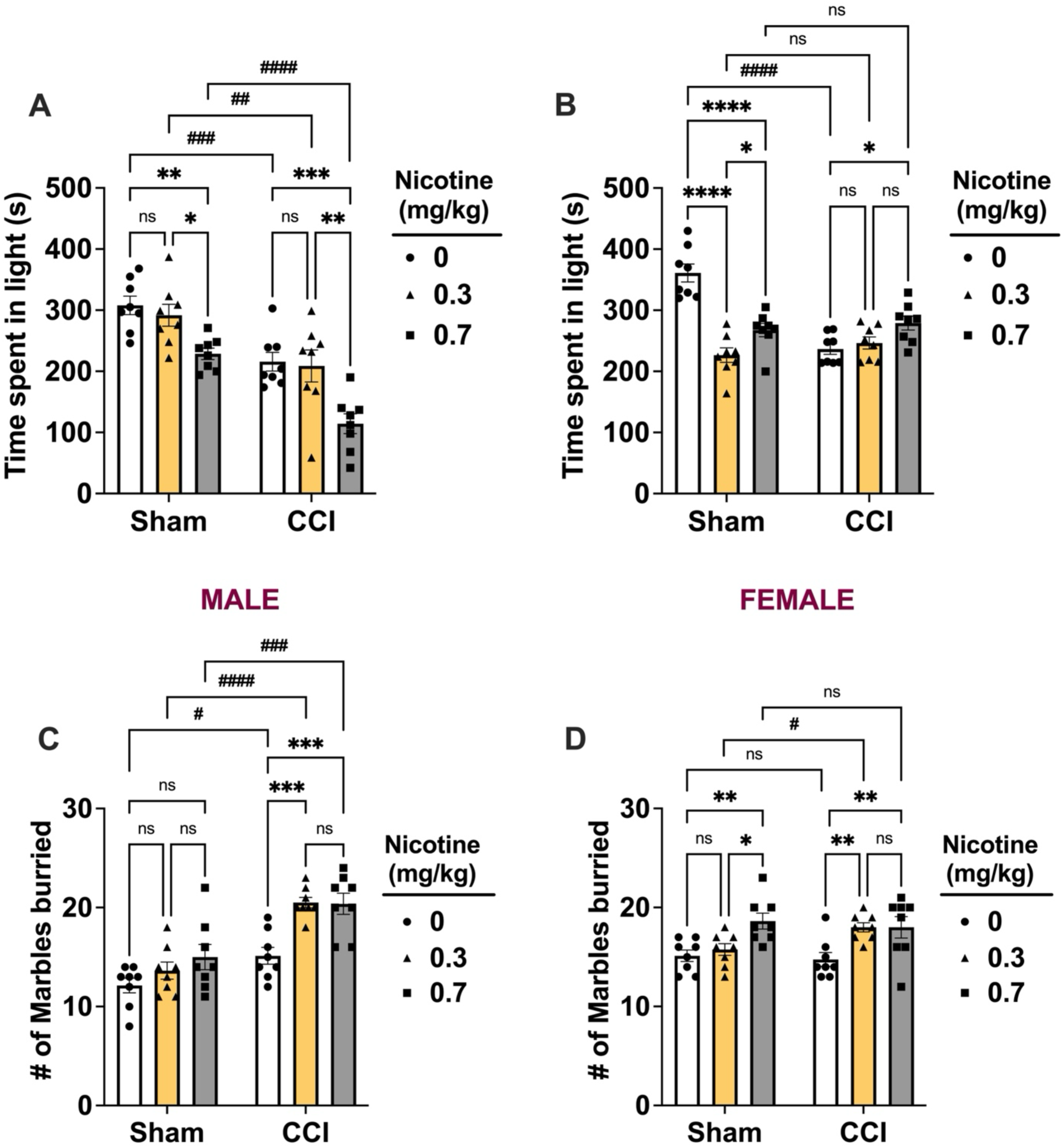
Impact of chronic pain on anxiety-like behaviors during nicotine withdrawal. Rats underwent sham or chronic constriction injury (CCI) surgery. Two weeks after surgery, animals received 14 days of subcutaneous injections of saline, 0.3 mg/kg, or 0.7 mg/kg nicotine. Approximately 24 h after the final injection, anxiety-like behavior was assessed using the light-dark transition test and marble burying test. Time spent in the light compartment was recorded for male (A) and female (B) rats, and the number of marbles buried was recorded for male (C) and female (D) rats. Data are expressed as mean ± SEM. \**p* < 0.05 compared with saline; #*p* < 0.05 compared with sham.

In females, nicotine decreased time spent in the light side only in sham rats (Figure 3B). A two-way ANOVA revealed significant main effects of nicotine [F_(2, 42)_ = 15.11, *p* < 0.0001], chronic pain [F_(1, 42)_ = 11.12, *p* = 0.0018], and interactions between nicotine and chronic pain [F_(2, 42)_ = 25.29, *p* < 0.0001] for female rats (Figure 3B). In female sham rats, both 0.3 and 0.7 mg/kg nicotine significantly decreased time spent in light compared with saline-treated rats, with 0.3 mg/kg nicotine had greater effects (*p* < 0.05). This suggests nicotine at both doses induced anxiety-like behavior but 0.3 had the highest impact. In female CCI rats, saline-treated rats showed a significant decrease in time spent in light compared with their sham controls, which suggests chronic pain induced anxiety-like behavior (*p* < 0.0001). While 0.3 mg/kg nicotine had no effect on time spent in light, 0.7 mg/kg nicotine significantly increased time spent in light compared with saline-treated CCI rats (*p* < 0.05), which indicates nicotine partially rescued pain-induced anxiety-like behavior.

Additional three-way ANOVA analyzed interactions between chronic pain state, nicotine dose, and sex effects (Supplementary Figure 2A). We found significant main effects for nicotine dose [F_(2, 21)_ = 20.35, *p* < 0.0001], sex [F_(1, 21)_ = 30.23, *p* < 0.0001], chronic pain [F_(1, 21)_ = 39.72, *p* < 0.0001], and global interactions between these three [F_(2, 21)_ = 8.470, *p* = 0.0020]. Additionally, there were significant interactions between nicotine dose and chronic pain state [F_(2, 21)_ = 5.166, *p* = 0.0150], between nicotine dose and sex [F_(2, 21)_ = 19.42, *p* < 0.0001], and between chronic pain and sex [F_(1, 21)_ = 14.93, *p* = 0.0009].

As a complementary measure of anxiety-like behavior, a marble burying test was also performed. Increased number of buried marbles is identified as anxiety-like behavior of withdrawal. In males, nicotine increased number of buried marbles in CCI rats but not in sham rats (Figure 3C). A two-way ANOVA revealed significant main effects of nicotine [F_(2, 42)_ = 11.34, *p* = 0.0001] and chronic pain [F_(1, 42)_ = 45.94, *p* < 0.0001], but no interactions between nicotine and chronic pain [F_(2, 42)_ = 0.2.262, *p* = 0.1166] for male rats (Figure 3C). Although a slight increase was observed, nicotine did not significantly increase on number of buried marbles in male sham rats (*p* > 0.05). In male CCI rats, saline-treated rats showed a significant increase on the number of buried marbles compared with their sham controls, which suggests chronic pain induced anxiety-like behavior (*p* < 0.05). Moreover, both 0.3 and 0.7 mg/kg significantly increased the number of buried marbles compared with saline-treated CCI rats without a dose relation (*p* < 0.001). This indicates nicotine induced anxiety-like behavior in male CCI rats. Post hoc analysis also revealed that the same nicotine dose increased the number of buried marbles in CCI rats compared to sham controls, indicating that chronic pain amplifies the anxiety-like behavior of nicotine withdrawal in males.

In females, saline-treated sham and CCI rats buried similar number of marbles, but nicotine increased the number of buried marbles both in sham and CCI rats (Figure 3D). A two-way ANOVA revealed significant main effects of nicotine [F_(2, 42)_ = 10.72, *p* = 0.0002], but no main effects for chronic pain [F_(1, 42)_ = 0.4868, *p* = 0.4892] or interactions between nicotine and chronic pain [F_(2, 42)_ = 2.371, *p* = 0.1058] for female rats (Figure 3D). In female sham rats, only 0.7 mg/kg nicotine significantly increased the number of buried marbles (*p* < 0.01), but both 0.3 and 0.7 mg/kg nicotine increased number of buried marbles in CCI rats (*p* < 0.01), which suggests nicotine withdrawal-induced anxiety-like behavior is more pronounced in the context of chronic pain.

Additional three-way ANOVA analyzed interactions between chronic pain state, nicotine dose, and sex effects (Supplementary Figure 2B). We found significant main effects for nicotine dose [F_(2, 21)_ = 13.25, *p* = 0.0002] and chronic pain [F_(1, 21)_ = 34.95, *p* < 0.0001], but no main effects of sex [F_(1, 21)_ = 1.964, *p* = 0.1757]. There was also no global interactions between these three [F_(2, 21)_ = 0.9058, *p* = 0.4194]. However, there were significant interactions between nicotine dose and chronic pain state [F_(2, 21)_ = 4.230, *p* = 0.0286] and between chronic pain and sex [F_(1, 21)_ = 34.33, *p* < 0.0001], but no interactions between nicotine dose and sex [F_(2, 21)_ = 1.085, *p* = 0.3562].

### 3.3. Impact of chronic pain on response to stress during nicotine withdrawal

To examine the impact of chronic pain on stress responsivity during nicotine withdrawal, we performed the forced swim test 72 h after the last injection of nicotine or saline. In male rats, a two-way ANOVA revealed significant main effects for chronic pain [F_(1, 42)_ = 11.82, *p* = 0.0013], but no main effects of nicotine [F_(2, 42)_ = 0.6720, *p* = 0.5161], or interactions between nicotine and chronic pain [F_(2, 42)_ = 0.8701, *p* = 0.4263 (Figure 4A). Planned comparisons revealed that chronic pain significantly increased the immobility time in saline-treated CCI rats compared with their corresponding sham controls (*p* < 0.01), and nicotine at any dose did not further alter immobility time either in sham rats or CCI rats (*p* > 0.05). However, immobility time in 0.7 mg/kg nicotine-treated CCI rats showed significantly higher duration of immobility compared with 0.7 mg/kg nicotine-treated sham rats (*p* < 0.05). In females, a two-way ANOVA revealed significant main effects of nicotine [F_(2, 42)_ = 4.837, *p* = 0.0129] and interactions between nicotine and chronic pain [F_(2, 42)_ = 4.195, *p* = 0.0218], but no main effects for chronic pain [F_(1, 42)_ = 0.8110, *p* = 0.3730] for female rats (Figure 4B). Saline-treated sham and CCI rats showed a similar duration of immobility time (*p* > 0.05). Nicotine at any doses did not alter immobility time (*p* > 0.05). However, 0.7 mg/kg nicotine-treated CCI rats significantly decreased the immobility time compared with their saline controls and corresponding sham controls (*p* < 0.05). This suggests nicotine dose-dependently decreased stress response in females under chronic pain condition.

**Figure 4.**
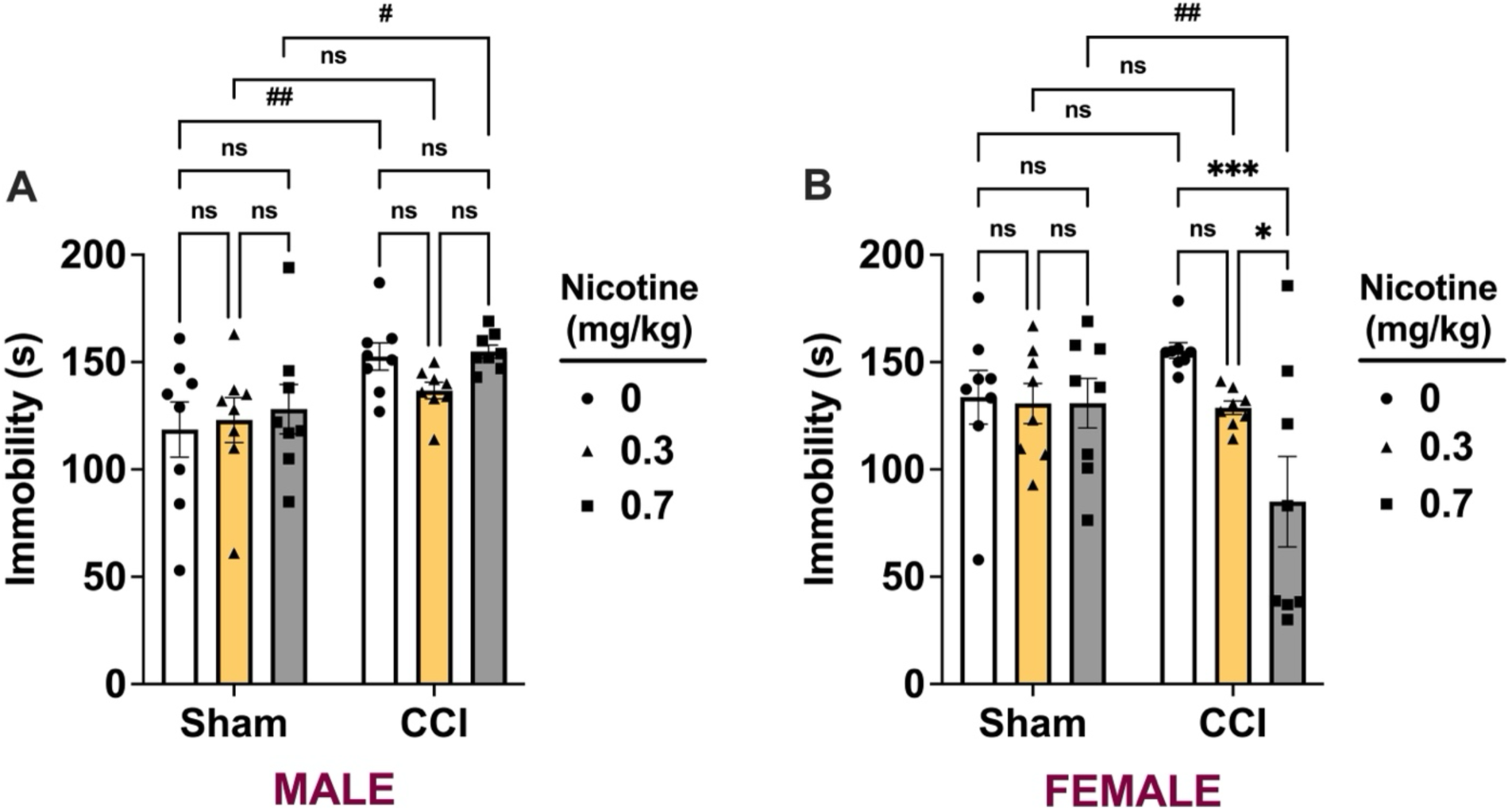
Impact of chronic pain on stress response during nicotine withdrawal. Rats underwent sham or chronic constriction injury (CCI) surgery. Two weeks after surgery, animals received 14 days of subcutaneous injections of saline, 0.3 mg/kg, or 0.7 mg/kg nicotine. Habituation to the forced swim test was conducted 48 h after the final injection, followed by testing at 72 h. Immobility time was recorded for male (A) and female (B) rats. Data are expressed as mean ± SEM. \**p* < 0.05 compared with saline; #*p* < 0.05 compared with sham.

Additional three-way ANOVA analyzed interactions between chronic pain state, nicotine dose, and sex effects (Supplementary Figure 3). No significant main effects were found for chronic pain [F_(1, 21)_ = 2.943, *p* = 0.1009], sex [F_(1, 21)_ = 2.035, *p* = 0.1684], and nicotine dose [F_(2, 21)_ = 2.243, *p* = 0.1309]. There was also no global interactions between these three [F_(2, 21)_ = 1.793, *p* = 0.1910]. However, there were significant interactions between nicotine dose and chronic pain state [F_(2, 21)_ = 5.300, *p* = 0.0137], and between chronic pain and sex [F_(1, 21)_ = 5.258, *p* = 0.0323], and interactions between nicotine dose and sex [F_(2, 21)_ = 4.967, *p* = 0.0171].

### 3.4. Impact of chronic pain on pain sensitivity during nicotine withdrawal

Mechanical sensitivity was assessed with the von Frey test on the nicotine withdrawal day. In male rats, a two-way ANOVA revealed significant main effects of chronic pain [F_(1, 42)_ = 53.62, *p* < 0.0001], nicotine [F_(2, 42)_ = 18.18, *p* < 0.0001], and interactions between nicotine and chronic pain [F_(2, 42)_ = 5.786, *p* = 0.0060 (Figure 5A). Chronic pain significantly decreased the paw withdrawal thresholds in saline-treated CCI rats compared with their corresponding sham controls (*p* < 0.0001). In sham rats, nicotine at both doses also decreased the paw withdrawal thresholds compared with saline treated rats (*p* < 0.0001). This suggests the development of mechanical allodynia in male sham rats during nicotine withdrawal. In CCI rats, nicotine further decreased paw withdrawal thresholds, but it did not reach a significant level (*p* > 0.05). The paw withdrawal thresholds in CCI rats were significantly less than their corresponding sham control rats (*p* < 0.05).

**Figure 5.**
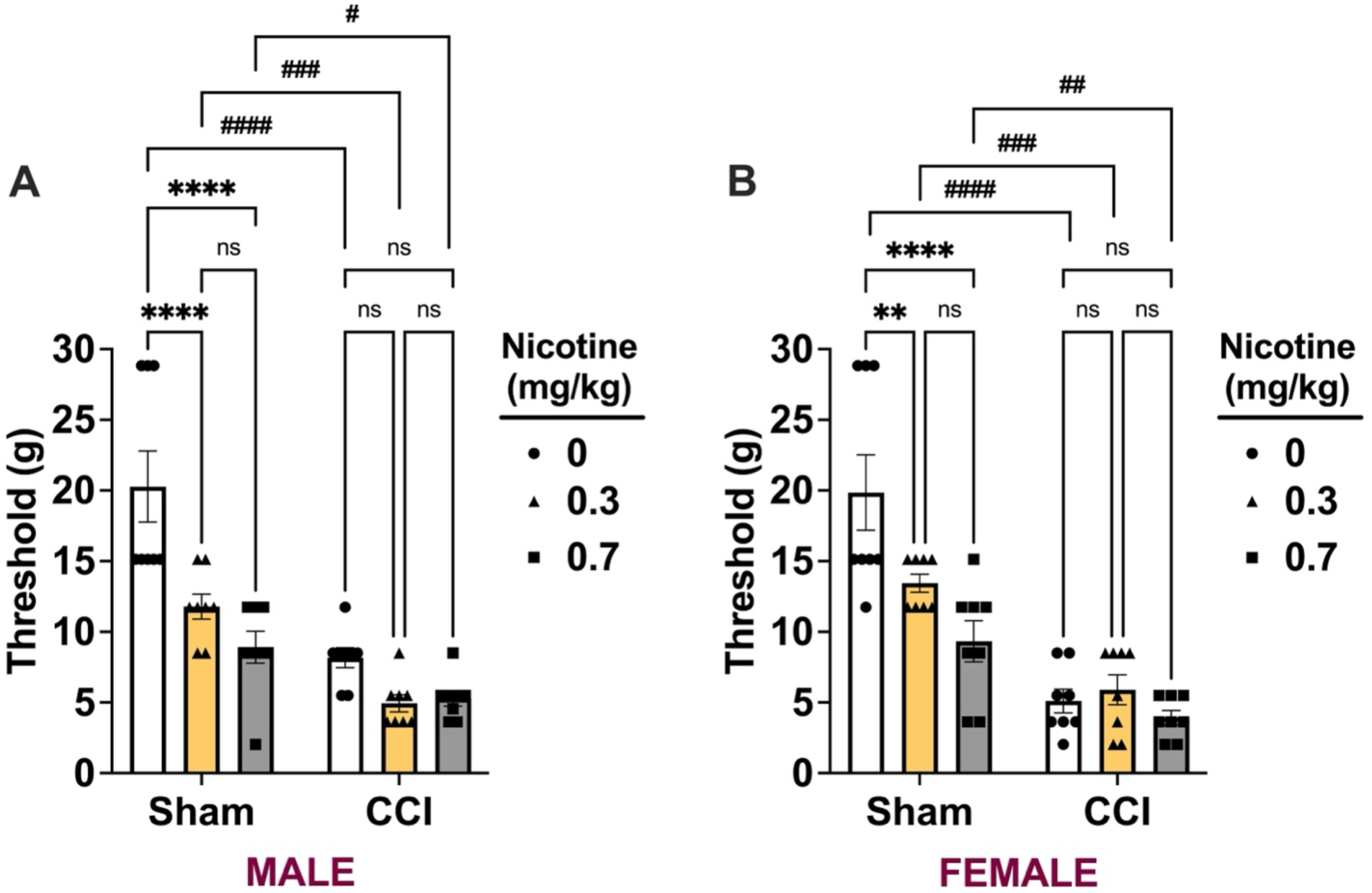
Impact of chronic pain on pain sensitivity during nicotine withdrawal. Rats underwent sham or chronic constriction injury (CCI) surgery. Two weeks after surgery, animals received 14 days of subcutaneous injections of saline, 0.3 mg/kg, or 0.7 mg/kg nicotine. Approximately 24 h after the final injection, pain sensitivity was assessed using the von Frey test. Paw withdrawal thresholds were recorded for male (A) and female (B) rats. Data are expressed as mean ± SEM. \**p* < 0.05 compared with saline; #*p* < 0.05 compared with sham.

In female rats, a two-way ANOVA revealed significant main effects of chronic pain [F_(1, 42)_ = 65.56, *p* < 0.0001], nicotine [F_(2, 42)_ = 8.801, *p* = 0.0006], and interactions between nicotine and chronic pain [F_(2, 42)_ = 6.142, *p* = 0.0046 (Figure 5B). Chronic pain significantly decreased the paw withdrawal thresholds in saline-treated CCI rats compared with their corresponding sham controls (*p* < 0.0001). In sham rats, nicotine at both doses also decreased the paw withdrawal thresholds compared with saline treated rats (*p* < 0.01). This suggests the development of mechanical allodynia in female sham rats during nicotine withdrawal. In CCI rats, nicotine at both doses did not change the paw withdrawal thresholds (*p* > 0.05). The paw withdrawal thresholds in CCI rats were significantly less than their corresponding sham control rats (*p* < 0.05).

Additional three-way ANOVA evaluated the interactions of chronic pain state, nicotine dose, and sex on paw withdrawal thresholds (Supplementary Figure 4). Significant main effects were observed for nicotine dose [F_(2, 21)_ = 28.12, *p* = 0.0001] and surgery [F_(1, 21)_ = 207.4, *p* < 0.0001], whereas sex had no significant effect [F_(1, 21)_ = 0.1104, *p* = 0.7430]. No global three-way interaction among surgery, dose, and sex was detected [F _(2, 21)_ = 0.1119, *p* = 0.8947]. A robust dose x surgery interaction indicated that the influence of nicotine dose differed between sham and CCI groups [F_(2, 21)_ = 20.58, *p* < 0.0001]. Neither dose x sex [F_(2, 21)_ = 0.9737, *p* = 0.3941] nor surgery x sex [F_(1, 21)_ = 1.028, *p* = 0.3221] interactions reached significance, indicating no evidence for sex-dependent modulation of mechanical sensitivity.

On the nicotine withdrawal day, both 0.3 and 0.7 mg/kg nicotine exposure decreased paw withdrawal thresholds compared with saline controls, indicating the development of mechanical allodynia in both males and females (*p* < 0.05). In sham rats, thresholds dropped from ∼20 g in saline groups (male sham: 20.3 g; female sham: 19.9 g) to ∼11-13 g with 0.3 mg/kg nicotine (male: 11.8 g; female: 13.4 g) and ∼9 g with 0.7 mg/kg nicotine (male: 8.9 g; female: 9.3 g). In CCI animals, thresholds were already reduced by surgery (male saline: 8.2 g; female saline: 5.1 g; *p* < 0.05) and remained low with nicotine exposure (0.3 mg/kg: 4.9 for males and 5.9 g for females; 0.7 mg/kg: 5.3 g for males and 3.9 g for females). At the 0.7 mg/kg nicotine dose, sham and CCI rats did not differ significantly on the withdrawal day (*p* > 0.05), and both groups showed robust allodynia.

#### Antinociceptive and antiallodynic effects of nicotine over time

On Day 0, nicotine (0.3 and 0.7 mg/kg) increased paw withdrawal thresholds in sham rats, indicating antinociceptive effects (Figure 6). In male rats, separate two-way RM ANOVA revealed significant main effects of nicotine dose and time, as well as a nicotine dose x time interaction on Days 0 and 7 [Day 0, F_nicotine dose (2, 21)_ = 30.95, *p* < 0.0001, F_time (2.783, 58.44)_ = 62.72, *p* < 0.0001, and F_nicotine dose and time (5.565, 58.44)_ = 19.17, *p* < 0.0001; Day 7, F_nicotine dose (2, 21)_ = 21.74, *p* < 0.0001, F_time (2.90, 60.90)_ = 57.23, *p* < 0.0001, and F_nicotine dose and time (5.80, 60.90)_ = 15.11, *p* < 0.0001, Figure 6A and B]. On Day 14, although other main effects remained significant, nicotine dose was no longer a significant main effect [Day 14, F_nicotine dose (2, 21)_ = 0.9457, *p* = 0.4043, F_time (1.695, 35.59)_ = 23.98, *p* < 0.0001], and F_nicotine dose and time (3.389, 35.59)_ = 5.787, *p* = 0.0018, Figure 6C]. In males, both doses produced antinociceptive effects, with 0.7 mg/kg showing a stronger and longer response that lasted up to 45 min. The effect appeared at 5 min, persisted through 30 min, and was diminished at 45 min (Figure 6A). On Day 7, the same pattern was observed, although the magnitude of the effect for 0.7 mg/kg was slightly reduced compared to Day 0 (Figure 6B). By Day 14, the antinociceptive effect was further reduced, with only a significant increase in withdrawal thresholds at 5 minutes (Figure 6C). Moreover, paw withdrawal thresholds significantly reduced at time 0 (before nicotine injection) compared to pre-surgery baseline values in both doses, indicating development of mechanical hypersensitivity.

**Figure 6.**
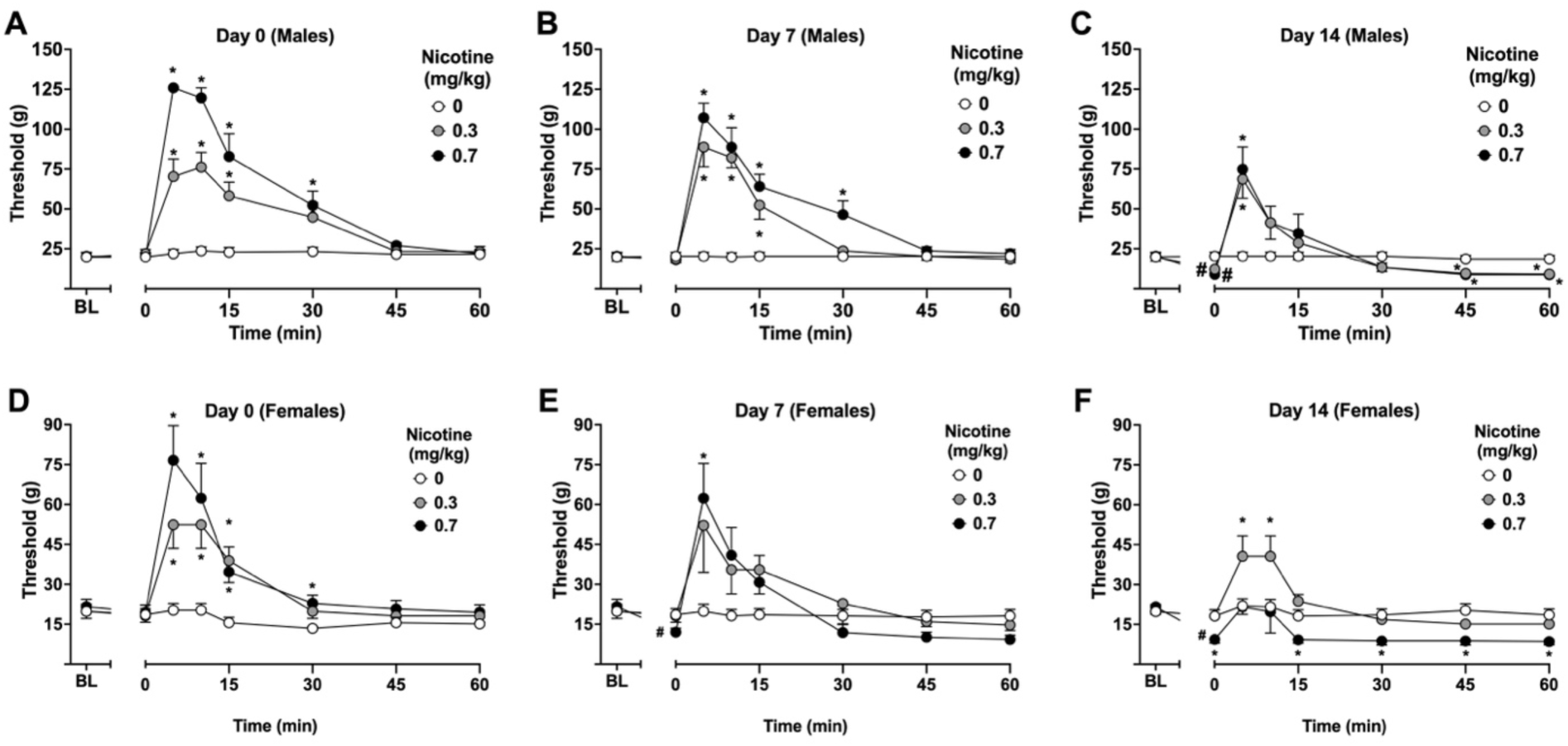
Antinociceptive effects of nicotine after acute and repeated exposure in sham rats. Rats underwent sham surgery. Paw withdrawal thresholds were assessed using the von Frey test at baseline (BL, Day −14) before surgery. Two weeks after surgery (Day 0), animals received subcutaneous injections of saline, 0.3 mg/kg, or 0.7 mg/kg nicotine. Testing was conducted on Day 0, Day 7, and Day 14. On each test day, thresholds were recorded immediately before injection (0 min) and at 5, 10, 15, 30, 45, and 60 min post-injection to assess the time course of nicotine’s acute (Day 0) and chronic (Days 7 and 14) antinociceptive effects. Panels show male rats (A, Day 0; B, Day 7; C, Day 14) and female rats (D, Day 0; E, Day 7; F, Day 14). Data are expressed as mean ± SEM. \**p* < 0.05 compared with saline. ^#^*p* < 0.05 compared with its baseline.

In female sham rats, separate two-way RM ANOVA revealed significant main effects of nicotine dose and time, as well as a nicotine dose x time interaction on Days 0 and 14 [Day 0, F_nicotine dose (2, 21)_ = 6.863, *p* = 0.0051, F_time (2.079, 43.67)_ = 22.39, *p* < 0.0001, and F_nicotine dose and time (4.159, 43.67)_ = 5.197, *p* < 0.0001; Day 14, F_nicotine dose (2, 21)_ = 5.853, *p* = 0.0095, F_time (1.940, 40.74)_ = 12.42, *p* < 0.0001, and F_nicotine dose and time (3.880, 40.74)_ = 3.610, *p* = 0.0138, Figure 6D and F]. On Day 7, although other main effects remained significant, nicotine dose was not a significant main effect [Day 7, F_nicotine dose (2, 21)_ = 0.9447, *p* = 0.4047, F_time (2.087, 43.82)_ = 13.29, *p* < 0.0001, and F_nicotine dose and time (4.174, 43.82)_ = 3.541, *p* = 0.0127, Figure 6E]. Like males, nicotine also produced dose-dependent antinociception on Day 0 in females. The antinociceptive effects were evident at 5 and lasted up to 45 min. (Figure 6D). On Day 7, the responses induced by 0.7 mg/kg nicotine were weaker, and nicotine at this dose also produced mechanical hypersensitivity at time 0 (Figure 6E). In females, the mechanical hypersensitivity started on day 7 and continued day 14. The antinociceptive effect was also minimal on Day 14; nicotine at 0.7 mg/kg showed lower thresholds compared to saline group, suggesting a loss of efficacy with repeated exposure and development of potential allodynia (Figure 6F).

CCI reduced paw withdrawal thresholds compared to baseline values in both male and female rats, indicating development of allodynia and this effect remained at all time 0 points (Figure 7). Nicotine (0.3 and 0.7 mg/kg) increased paw withdrawal thresholds, indicating antiallodynic effects (Figure 7). In male CCI rats, separate two-way RM ANOVA revealed significant main effects of nicotine dose and time, as well as a nicotine dose x time interaction on Days 0 and 7 [Day 0, F_nicotine dose (2, 21)_ = 32.62, *p* < 0.0001, F_time (2.676, 56.20)_ = 40.97, *p* < 0.0001, and F_nicotine dose and time (5.352, 56.20)_ = 14.29, *p* < 0.0001; Day 7, F_nicotine dose (2, 21)_ = 12.73, *p* = 0.0002], F_time (1.874, 39.35)_ = 29.54, *p* < 0.0001, and F_nicotine dose and time (3.748, 39.35)_ = 11.65, *p* < 0.0001, Figure 7A and B]. On Day 14, although other main effects remained significant, nicotine dose was no longer a significant main effect [Day 14, F_nicotine dose (2, 21)_ = 2.662, *p* = 0.0932, F_time (1.837, 38.58)_ = 14.96, *p* < 0.0001], and F_nicotine dose and time (3.675, 38.58)_ = 4.364, *p* = 0.0063, Figure 7C]. In males, both doses produced antiallodynic effects, with 0.7 mg/kg showing a stronger response that lasted up to 30 min. The effect appeared at 5 min, persisted through 15 min, and was diminished at 30 min (Figure 7A). On Day 7, nicotine again increased thresholds in CCI male rats. However, the antiallodynic effects were slightly reduced at both doses compared to Day 0 (Figure 7B). By Day 14, these effects were further diminished, with only the 0.7 mg/kg dose remaining effective at 5 min (Figure 7C).

**Figure 7.**
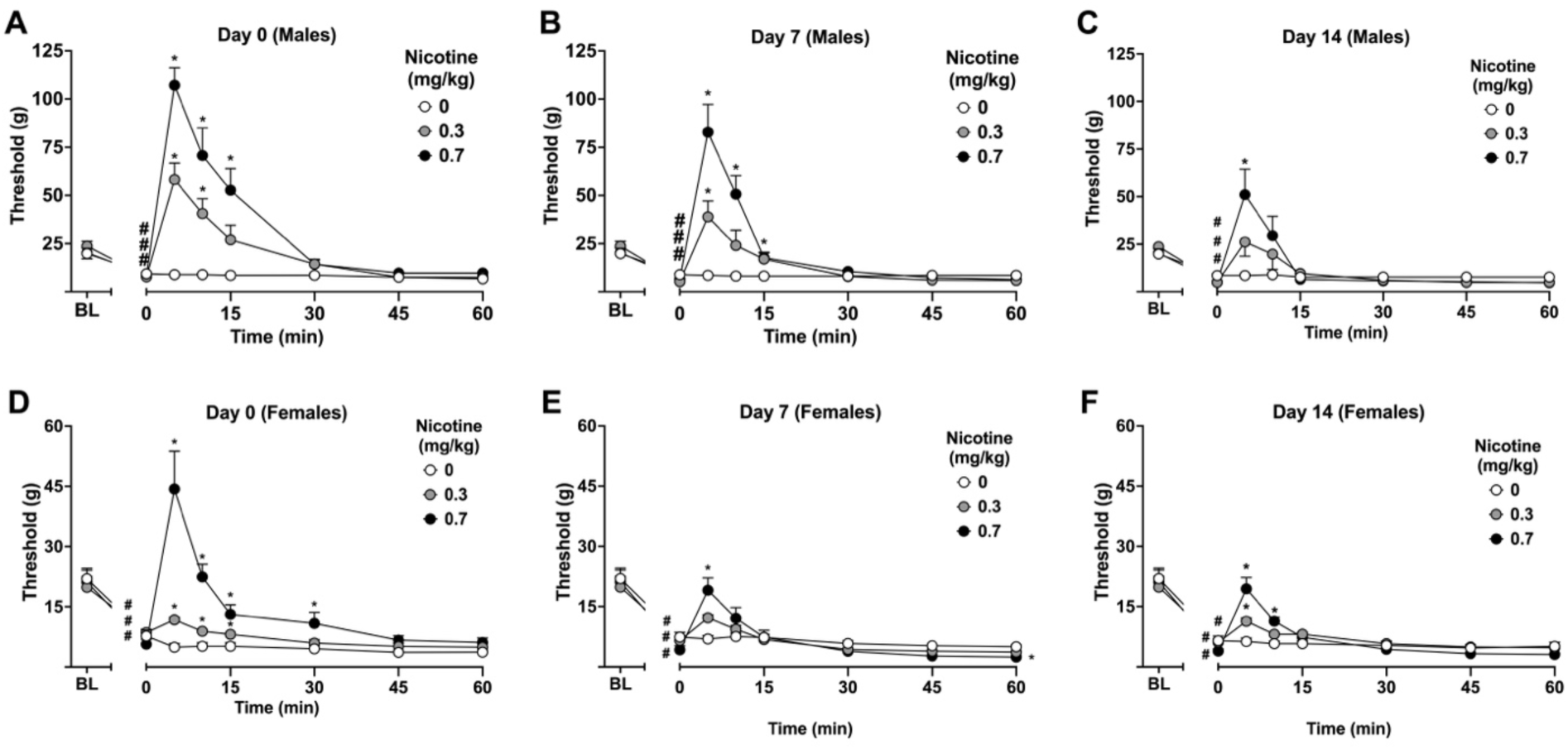
Antiallodynic effects of nicotine after acute and repeated exposure in rats with chronic pain. Rats underwent chronic constriction injury (CCI) surgery. Paw withdrawal thresholds were assessed using the von Frey test at baseline (BL, Day −14) before surgery. Two weeks after surgery (Day 0), animals received subcutaneous injections of saline, 0.3 mg/kg, or 0.7 mg/kg nicotine. Testing was conducted on Day 0, Day 7, and Day 14. On each test day, thresholds were recorded immediately before injection (0 min) and at 5, 10, 15, 30, 45, and 60 min post-injection to assess the time course of nicotine’s acute (Day 0) and chronic (Days 7 and 14) antiallodynic effects. Panels show male rats (A, Day 0; B, Day 7; C, Day 14) and female rats (D, Day 0; E, Day 7; F, Day 14). Data are expressed as mean ± SEM. \**p* < 0.05 compared with saline. #*p* < 0.05 compared with its baseline.

In female CCI rats, separate two-way RM ANOVA revealed significant main effects of nicotine dose and time, as well as a nicotine dose x time interaction on Day 0 [Day 0, F_nicotine dose (2, 21)_ = 24.69, *p* < 0.0001, F_time (1.954, 41.04)_ = 21.05, *p* < 0.0001, and F_nicotine dose and time (3.908, 41.04)_ = 8.960, *p* < 0.0001, Figure 7D]. On Days 7 and 14, although other main effects remained significant, nicotine dose was no longer a significant main effect [Day 7, F_nicotine dose (2, 21)_ = 0.3043, *p* = 0.7408, F_time (2.335, 49.03)_ = 54.53, *p* < 0.0001, and F_nicotine dose and time (4.669, 49.03)_ = 3.694, *p* = 0.0075; Day 14, F_nicotine dose (2, 21)_ = 1.171, *p* = 0.3295, F_time (2.105, 44.22)_ = 68.58, *p* < 0.0001, and F_nicotine dose and time (4.211, 44.22)_ = 68.58, *p* = 0.0009, Figure 7E and F]. In females, nicotine (0.3 and 0.7 mg/kg) also induced antiallodynic effects, although the magnitude was markedly lower than in males. Both doses produced modest increases in paw withdrawal thresholds, with the 0.7 mg/kg dose showing a slightly stronger and more consistent effect on Day 0 (Figure 7D). The antiallodynic response started at 5 min, persisted to 15 min for 0.3 mg/kg and to 30 min for 0.7 mg/kg, and was diminished by 45 min (Figure 7D). On Day 7, nicotine produced antiallodynic effects only at the 0.7 mg/kg dose (Figure 7E). By Day 14, these effects were minimal, with only a transient but significant response observed, suggesting reduced efficacy with repeated exposure (Figure 7F).

To compare the effects of nicotine between sham and CCI conditions across sexes, we focused on the 5 min time point, where nicotine produced peak analgesic-like effects, reflected as antinociception in sham rats and antiallodynia in CCI rats in Supplementary Figure 5. In male rats, a two-way RM ANOVA revealed significant main effects of nicotine [F_(5, 126)_ = 53.71, *p* < 0.0001], time [F_(2, 126)_ = 11.59, *p* < 0.0001], and interactions between nicotine and time [F_(10, 126)_ = 2.496, *p* = 0.0091 (Supplementary Figure 5A). In sham males, nicotine (0.3 and 0.7 mg/kg) induced significant antinociceptive effects at 5 min on Day 0 (*p* < 0.05), with a stronger response at 0.7 mg/kg. These effects remained significant on Day 7, although reduced in magnitude, particularly at 0.7 mg/kg, and were further diminished by Day 14, where the response at 0.7 mg/kg was significantly lower than Day 0 (*p* < 0.05). In CCI males, nicotine produced significant antiallodynic effects at both doses on Day 0, with a stronger response at 0.7 mg/kg. However, the magnitude of these effects was lower than in sham and declined over time, with reduced responses on Day 7 and further attenuation by Day 14 at both doses relative to Day 0 (*p* < 0.05). In female rats, a two-way RM ANOVA revealed significant main effects of nicotine [F_(5, 126)_ = 21.69, *p* < 0.0001], time [F_(2, 126)_ = 6.457, *p* = 0.0021], and interactions between nicotine and time [F_(10, 126)_ = 2.876, *p* = 0.0029 (Supplementary Figure 5B). In sham females, nicotine (0.3 and 0.7 mg/kg) induced antinociceptive effects at 5 min on Day 0 (*p* < 0.05), although the magnitude was lower than in males. These effects were maintained on Day 7 (*p* < 0.05) but were absent by Day 14 (*p* > 0.05), with the response at 0.7 mg/kg significantly reduced relative to Day 0 (*p* < 0.05). In CCI females, nicotine produced a limited antiallodynic effect on Day 0, observed only at the 0.7 mg/kg dose (*p* < 0.05). Across Days 7 and 14, these effects were minimal, with no evidence of sustained or significant antiallodynic responses.

For Supplementary Figure 5, an additional three-way repeated-measures ANOVA evaluated the interactions between chronic pain state, nicotine dose, and sex, with time (day) included as a within-subject factor on paw withdrawal threshold at 5 minutes. Significant main effects were observed for chronic pain state [F_(1, 84)_ = 72.13, p < 0.0001], sex [F_(1, 84)_ = 87.75, *p* < 0.0001], nicotine dose [F_(2, 84)_ = 109.93, *p* < 0.0001], and time [F_(2, 168)_ = 19.52, *p* < 0.0001]. The interaction between chronic pain state and nicotine dose was significant [F_(2, 84)_ = 5.02, *p* = 0.0087], as was the nicotine dose x time interaction [F_(4, 168)_ = 10.35, *p* < 0.0001], indicating that the effects of nicotine dose varied over time. There was also a significant sex x nicotine dose interaction [F_(2, 84)_ = 24.8478, *p* < 0.0001]. No significant interactions were detected for chronic pain state x sex [F_(1, 84)_ = 0.21, *p* = 0.651], chronic pain state x sex x nicotine dose [F_(2, 84)_ = 0.009, *p* = 0.991], chronic pain state x time [F_(2, 168)_ = 1.36, *p* = 0.260], sex x time [F_(2, 168)_ = 1.19, *p* = 0.306], chronic pain state x sex x time [F_(2, 168)_ = 2.01, *p* = 0.137], chronic pain state x nicotine dose x time [F_(4, 168)_ = 0.99, *p* = 0.417], sex x nicotine dose x time [F_(4, 168)_ = 0.20, *p* = 0.940], or the four-way interaction among chronic pain state, sex, nicotine dose, and time [F_(4, 168)_ = 1.18, *p* = 0.323].

#### Effects of surgery and chronic nicotine exposure on mechanical sensitivity

Weekly paw withdrawal thresholds were tracked in males and females from baseline Post Surgery Day 0 through Post Surgery Day 35 (Figure 8). After baseline assessment, animals underwent either sham or CCI surgery. On Day 14, following threshold testing, daily injections of saline or nicotine (0.3 or 0.7 mg/kg) were initiated and continued for 14 days. Treatment was then discontinued, and thresholds were measured again one week later on Day 35.

**Figure 8.**
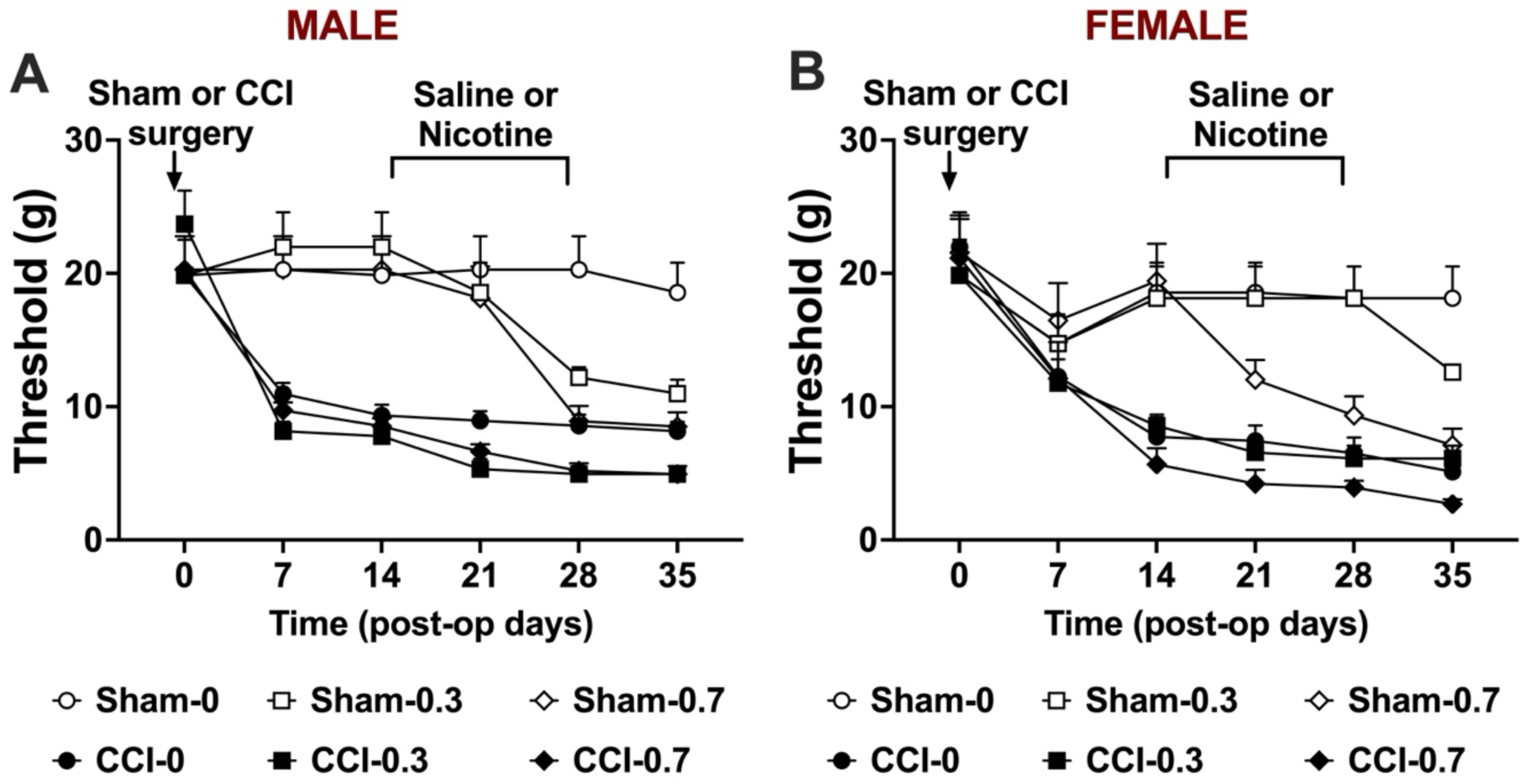
Impact of pain induction and chronic nicotine exposure on paw withdrawal thresholds. Weekly paw withdrawal thresholds (g) were measured in male (A) and female (B) rats from baseline (Day 0) through post-op Day 35. Following baseline testing, animals underwent sham or chronic constriction injury (CCI) surgery. On post-op Day 14, after threshold assessment, animals received daily subcutaneous injections of saline or nicotine (0.3 or 0.7 mg/kg) for 14 days, after which injections were discontinued. Animals were maintained for an additional week, and final paw withdrawal thresholds were assessed on post-op Day 35. Data are expressed as mean ± SEM.

In male rats, a three-way ANOVA evaluated the interactions between chronic pain state, and nicotine dose, with time included as a within-subject factor (Figure 8A). Significant main effects were observed for chronic pain state [F_(1, 42)_ = 62.32, *p* < 0.0001], and time [F_(5, 210)_ = 48.46, *p* < 0.0001], with a modest trend for nicotine dose [F_(2, 42)_ = 2.64, *p* = 0.083]. There was no significant chronic pain state x nicotine dose interaction [F_(2, 42)_ = 0.36, p = 0.697]. Several interactions involving time were significant, indicating that the effects of surgery and nicotine dose changed across sessions: chronic pain state x time [F_(5, 210)_ = 18.97, p < 0.0001] and nicotine dose x time [F_(10, 210)_ = 3.54, *p* = 0.0031]. The chronic pain x nicotine dose x time interaction was nonsignificant [F_(10, 210_) = 1.72, p = 0.121]. In female rats, a three-way repeated-measures ANOVA evaluated the effects of chronic pain state and nicotine dose, with timepoint included as a within-subject factor (Figure 8B). Significant main effects were observed for chronic pain state [F_(1, 42)_ = 33.02, *p* < 0.0001] and time [F_(5, 210)_ = 58.30, *p* < 0.0001], whereas the main nicotine dose was nonsignificant [F_(2, 42)_ = 1.88, *p* = 0.165]. No significant chronic pain state x nicotine dose interaction was detected [F_(2, 42)_ = 0.19, *p* = 0.832]. Several interactions involving timepoint were significant, indicating that the effects of chronic pain state and nicotine dose changed across sessions: chronic pain state x time [F_(5, 210)_ = 17.74, *p* < 0.0001], nicotine dose x time [F_(10, 210)_ = 3.80, *p* = 0.00178], and chronic pain state x nicotine dose x time [F_(10, 210)_ = 2.22, *p* = 0.0486].

At baseline (Day 0), paw withdrawal thresholds were ∼19-23 g in both males (Figure 8A) and females (Figure 8B) across all groups. CCI surgery induced a significant reduction in paw withdrawal thresholds beginning on Day 7 and reaching a stable decrease by Day 14 compared with baseline values (*p* < 0.05), consistent with the development of mechanical allodynia. By Day 14, CCI surgery reduced thresholds to ∼8-9 g in males (Figure 8A) and ∼6-8 g in females (Figure 8B), while sham groups remained near baseline. A modest postoperative threshold decrease was observed in female sham rats, but this did not reach statistical significance. Paw withdrawal thresholds in sham saline and CCI saline groups remained stable between Days 14 and 35, indicating that repeated von Frey testing did not induce mechanical hypersensitivity (Figure 8).

Following initiation of nicotine treatment, female sham rats receiving 0.7 mg/kg nicotine showed a reduction in thresholds by Day 21, indicating nicotine-induced mechanical hypersensitivity (Figure 8B). By Day 28, both 0.3 and 0.7 mg/kg nicotine produced decreases in thresholds in male sham rats (Figure 8A), while 0.7 mg/kg continued to reduce thresholds in female sham rats (Figure 8B). Although Days 21 and 28 were classified as treatment days, the paw withdrawal threshold assessments occurred in the morning following the previous evening’s nicotine injection, meaning animals were likely in an early withdrawal state during testing. On Day 35, allodynia persisted in male sham rats treated with 0.3 or 0.7 mg/kg nicotine (Figure 8A) and in female sham rats treated with 0.7 mg/kg, with 0.3 mg/kg producing a modest additional decrease in female sham rats (Figure 8B).

In CCI animals treated with nicotine, reductions in paw withdrawal thresholds were observed at Days 21, 28, and 35 in males receiving 0.3 or 0.7 mg/kg (Figure 8A), and in females receiving 0.7 mg/kg (Figure 8B). However, these decreases were less pronounced than those observed in sham groups, suggesting that the impact of nicotine-induced hypersensitivity was blunted in the presence of pre-existing neuropathic pain or a floor effect for mechanical hypersensitivity in these rats.

By Day 35, sham saline animals maintained stable thresholds (∼18-19 g), but sham rats treated with 0.7 mg/kg nicotine showed marked reductions, reaching ∼8.5 g in males (Figure 8A) and ∼7.1 g in females (Figure 8B). CCI animals remained hypersensitive throughout, with males stabilizing around ∼4.9 g (Figure 8A) and females dropping as low as ∼2.7 g with 0.7 mg/kg nicotine (Figure 8B).

For Figure 8, an additional three-way repeated-measures ANOVA evaluated the interactions between chronic pain state, nicotine dose, and sex, with time included as a within-subject factor. Significant main effects were observed for chronic pain state [F_(1, 84)_ = 89.82, p < 0.0001], nicotine dose [F_(2, 84)_ = 4.07, p = 0.0205], and time [F_(5, 420)_ = 104.82, p < 0.0001], whereas sex had no significant effect [F_(1, 84)_ = 1.23, p = 0.270]. No significant three-way interaction among surgery, dose, and sex was detected [F_(2, 84)_ = 0.0091, p = 0.991]. Several interactions with time were observed. There were significant surgery x time [F_(5, 420)_ = 31.65, p < 0.0001] and nicotine dose x time interactions [F_(10, 420)_ = 5.61, p < 0.0001], indicating that both chronic pain state and nicotine dose differentially influenced paw withdrawal thresholds over time. A significant surgery x sex x time interaction [F_(5, 420)_ = 5.13, p = 0.0016] further suggested sex-dependent temporal effects of chronic pain. Additionally, a surgery x nicotine dose x time interaction was observed [F_(10, 420)_ = 2.96, p = 0.0075], indicating that the effects of nicotine dose on mechanical sensitivity over time differed between sham and CCI groups. A modest trend towards interaction was detected for sex x nicotine dose x time [F_(10, 420)_ = 1.75, p = 0.105], whereas the four-way interaction among surgery, sex, nicotine dose, and time was not significant [F_(10, 420)_ = 0.96, p = 0.458].

## 4. Discussion

Our findings provide the first preclinical evidence that ongoing chronic pain exacerbates nicotine withdrawal severity in a nicotine concentration- and sex-dependent manner, enhancing both physical and affective withdrawal symptoms. Repeated nicotine exposure led to tolerance to its analgesic-like effects and increased pain sensitivity that persisted after nicotine abstinence.

In humans, aversive abstinence syndrome typically emerges within 4 to 24 h after cessation of chronic nicotine exposure, and it gradually diminishes over the following 3 to 4 weeks [38]. Consistent with this time course, preclinical studies have shown that repeated nicotine exposure produces both somatic and affective signs of withdrawal during abstinence [19]. While mecamylamine-precipitated withdrawal produces rapid, robust signs via nicotinic receptor blockade, spontaneous withdrawal better models abstinence after repeated exposure [21]. In the present study, we used spontaneous withdrawal to assess a natural course of withdrawal symptoms.

In this study, 24 h nicotine abstinence induced physical withdrawal signs in a dose-dependent manner (Figure 2), consistent with prior preclinical nicotine studies [19]. Chronic pain further amplified nicotine withdrawal-associated physical withdrawal signs. As this is the first study to examine the impact of chronic pain on nicotine withdrawal, direct comparisons with prior preclinical studies are limited. However, our results are consistent with recent clinical consensus statement suggesting that chronic pain is associated with greater nicotine withdrawal severity in humans [28]. In human, it is difficult to determine whether chronic pain or nicotine dependence develops first. Our findings provide new insight into this question, as we directly examined the contribution of chronic pain to the severity of nicotine withdrawal. Sex differences were also observed in how chronic pain enhanced somatic signs, with female rats exhibiting higher withdrawal severity at the higher nicotine dose. Previous human studies reported that women are more sensitive to nicotine concentration and dependence than men [45,47,49,50,58]. In addition, women have lower cessation success rates [18,46,50,52]. These findings suggest that sex differences in nicotine sensitivity interact with chronic pain, influencing withdrawal severity, particularly at higher nicotine exposure.

The light-dark box is widely used to assess anxiety-like behavior, and prior studies show that nicotine withdrawal induces anxiety-like responses [2,64]. Consistent with this literature, nicotine withdrawal induced anxiety-like behavior in male rats in a dose-dependent manner in both sham and CCI groups (Figure 3A). Chronic pain alone also induced anxiety-like behavior in males, and nicotine withdrawal further exacerbated this effect, revealing that nicotine withdrawal-associated anxiety is enhanced in the presence of chronic pain. In female rats, nicotine withdrawal induced anxiety-like behavior in sham animals, and chronic pain alone also induced anxiety-like behavior (Figure 3B). However, under chronic pain conditions, nicotine at the higher dose reduced anxiety-like behavior rather than further increasing it. This indicates a sex-dependent modulation of anxiety-like responses to withdrawal in the context of chronic pain. Consistent with reports that women more often use nicotine for mood regulation [51], our findings support the notion that nicotine use in females may reflect self-medication strategy to reduce anxiety during withdrawal.

The marble burying test provided complementary insight into anxiety-like behavior during nicotine withdrawal (Figure 3 C and D). In males, nicotine exposure alone did not alter baseline anxiety-like behavior in sham rats, whereas chronic pain induced anxiety-like behavior on its own and increased anxiety-like responses during withdrawal. In females, baseline anxiety-like behavior was not altered by chronic pain alone; however, withdrawal increased marble burying behavior, with effects observed at the higher nicotine dose in sham rats and at both nicotine doses in CCI rats. These findings indicate that anxiety-like behavior measured by marble burying is differentially influenced by withdrawal and chronic pain in a sex-dependent manner, with males showing strong pain-driven effects and females showing greater sensitivity to nicotine withdrawal. The finding chronic pain alone induced anxiety-like behavior is consistent with prior studies showing elevated negative affect across multiple different assays [22,31,62,68] and contrary to the studies which did not show [27], and extends this work by demonstrating that chronic pain-induced anxiety-like behavior amplifies withdrawal-associated negative affect.

Nicotine withdrawal increases immobility (behavioral despair) in the forced swim test in rodents [20,56,69]. In this study, chronic nicotine exposure at either dose did not alter immobility in sham rats (Figure 4). Given that immobility was measured 72 hours after the final nicotine injection, it is possible that this observation window is outside the peak withdrawal time point for rats these nicotine dosages. Further, a precipitated withdrawal approach may be more sensitive for detecting effect in the forced swim test, as withdrawal-related depressive-like behavior in rats may occur within a relatively brief time window compared with mice [19]. Particularly, nicotine dose and duration of exposure can determine the severity of dependence in rodents [21]. However, chronic pain alone increased depressive-like behavior in males. This finding is consistent with previous reports showing that chronic pain increases immobility in male rodents [9,31,68], as well as with human reports suggesting depression in the context of chronic pain [33,55]. Male CCI rats treated with the higher nicotine dose also exhibited increased immobility compared with sham rats receiving the same dose, suggesting that depressive-like behavior in males was primarily driven by the chronic pain state rather than withdrawal itself.

In females, chronic pain alone produced only a slight, non-significant increase in immobility. However, nicotine exposure reduced immobility in female CCI rats in a dose-dependent manner. Based on clinical evidence showing higher rates of depression in women and the role of stress in nicotine use [5,15,36,63], we initially hypothesized that withdrawal would exacerbate depressive-like behavior in female CCI rats. In contrast, nicotine exposure attenuated depressive-like behavior in females under chronic pain. However, stress has been proposed as a key driver of nicotine use and dependence in women [63]. Additionally, nicotine can produce antidepressant effects [4,37,54,57], potentially contributing to the maintenance of nicotine use. Our sex-specific finding suggests an interaction between chronic pain and nicotine exposure that differs between males and females and raises the possibility that nicotine use in females may be partly maintained by its ability to reduce pain-associated depressive-like symptoms.

Nicotine withdrawal is associated with increased pain sensitivity [6,21,71]. In sham rats, paw withdrawal thresholds decreased during abstinence, consistent with withdrawal-associated pain sensitivity (Figure 4). In CCI rats, thresholds also decreased, particularly in males, but did not reach significance, likely due to a floor effect. While much of the previous literature on nicotine withdrawal-associated pain has focused on hyperalgesia [6,21], defined as increased responses to normally painful stimuli, the present findings extend this work by demonstrating allodynia, reflecting pain responses to normally innocuous stimuli during nicotine abstinence. Our allodynic outcomes are consistent with other studies that measured this outcome [71]. Thus, our results align with prior reports showing that nicotine withdrawal enhances pain sensitivity, while highlighting allodynia as an additional withdrawal-related pain outcome.

Nicotine can produce antinociceptive, antiallodynic, and antihyperalgesic effects across multiple preclinical pain models [3,7,16,30,65,71]. However, repeated nicotine exposure leads to tolerance to these effects [3,26,41,71]. As mentioned above, withdrawal from nicotine increases pain sensitivity [6,21,71]. This may reflect the heightened pain sensitivity seen in humans during early abstinence from nicotine use. Consistent with the literature, nicotine produced robust antinociceptive effects in sham rats in a concentration-, sex-, and time-dependent manner (Figure 6). These effects declined with repeated exposure, particularly at the higher dose, indicating tolerance, with greater sensitivity in females. Repeated nicotine exposure also lowered baseline paw withdrawal thresholds, indicating increased pain sensitivity. Overall, chronic nicotine exposure resulted in loss of antinociception and the development of hypersensitivity, consistent with prior reports. In line with previous report [7], nicotine produced anti-allodynic effects in chronic pain, which were concentration- and sex-dependent and persisted with repeated exposure, though with sex differences (Figure 7). In males, nicotine attenuated allodynia at both doses, with declining efficacy over time. In females, anti-allodynic effects were dose-dependent and sustained only at the higher dose. These findings indicate the development of tolerance, with greater dose dependence in females.

Nicotine has complex effects on pain as discussed above. Chronic nicotine use is a risk factor for the development and worsening of chronic pain conditions [59,60]. In contrast, acute nicotine reverses paclitaxel-induced neuropathic pain, while chronic exposure prevents its development in mice [30]. Given these opposing effects, we assessed weekly pain thresholds throughout the study (Figure 8). Repeated nicotine exposure induced mechanical hypersensitivity that persisted beyond administration. Prior studies show that chronic nicotine initially produces antinociception but later induces hypersensitivity [71] and neuropathic pain-like outcomes [61]. Our findings are consistent with these reports and extend these works by demonstrating prolonged, sex-dependent effects.

This study reveals that chronic pain exacerbates nicotine withdrawal severity in a concentration- and sex-dependent manner and that repeated nicotine exposure leads to tolerance to its analgesic-like effects. In pain-naive rats, repeated nicotine exposure induced mechanical hypersensitivity during withdrawal, supporting a reciprocal relationship between pain and nicotine use. A limitation is that the estrous cycle was not monitored in female rats, so the influence of ovarian hormones, which modulate nicotine-related behaviors including withdrawal [25], cannot be determined. Further studies are needed to test ovarian hormone contributions on this comorbidity.

## Acknowledgement

The authors thank Eve Andersen for her assistance with R analysis. Illustrations in Figure 1 were created with BioRender.com and sublicensed to the authors for use in publications.

## Ethics approval

Animal experiments were conducted in accordance with The Yale University Institutional Animal Care and Use Committee (IACUC) guideline (IACUC Protocol no. #2025-20661)

## Data Availability

The data underlying this study will be made available upon reasonable request to the corresponding author.

## Declaration of AI-assisted technologies

ChatGPT (OpenAI, GPT-5.3; Plus subscription) was used to refine wording and shorten text for clarity and concision at minimum level. No primary data or confidential information were provided to the tool. All content was reviewed by the authors, who take full responsibility for the accuracy and integrity of the manuscript.

## CRediT authorship contribution statement

Brianna Graham: Writing – original draft, review & editing, Data curation. Tyler Nelson & Sheela Tavakoli – review & editing, Data curation. Laura O’Dell – review & editing, Supervision. Nii A. Addy: Writing – review & editing, Supervision, Resources. Deniz Bagdas: Writing – original draft, review & editing, Supervision, Conceptualization, Methodology, Formal analysis, Data curation, Resources, Funding acquisition.

