## Supplementary for "Chronic pain exacerbates nicotine withdrawal severity in a sex-specific and dose-dependent manner"

**Supplementary Material.**

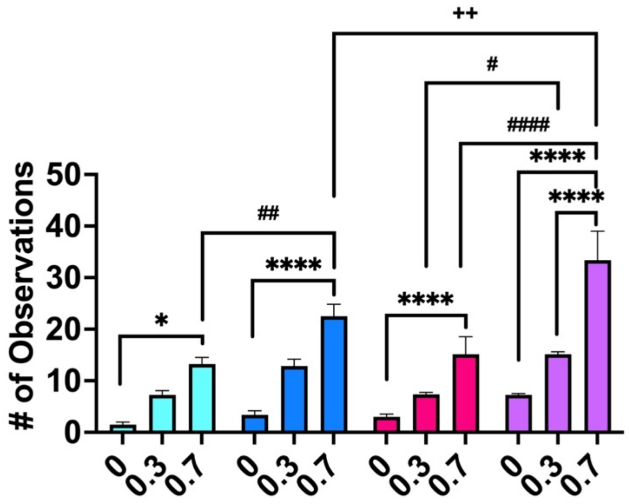


**Supplementary Figure 1: Sex differences in physical symptoms of nicotine withdrawal: three-way ANOVA.** A three-way ANOVA analyzed interactions between chronic pain state, nicotine dose, and sex effects. We found significant main effects for nicotine dose [$F_{(2, 21)} = 29.92$, $p < 0.0001$], sex [$F_{(1, 21)} = 8.202$, $p = 0.0093$], and chronic pain [$F_{(1, 21)} = 132.6$, $p < 0.0001$] but no global interactions between these three [$F_{(2, 21)} = 1.596$, $p = 0.2264$]. There were significant interactions between nicotine dose and chronic pain state [$F_{(2, 21)} = 21.28$, $p < 0.0001$] but no interactions between nicotine dose and sex [$F_{(2, 21)} = 0.669$, $p = 0.2125$]. However, there was significant interactions between chronic pain and sex [$F_{(1, 21)} = 6.376$, $p = 0.0197$], which revealed sex-dependent effects in chronic pain conditions.

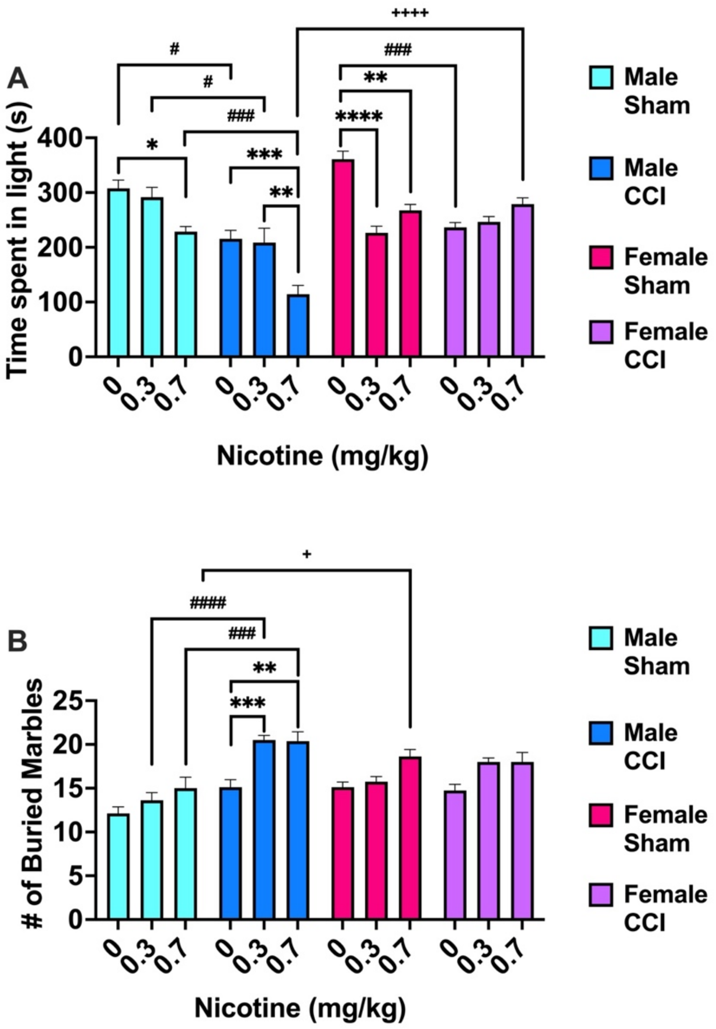


**Supplementary Figure 2: Sex differences in anxiety-like symptoms of nicotine withdrawal: three-way ANOVA.** Athree-way ANOVA analyzed interactions between chronic pain state, nicotine dose, and sex effects (Supplementary Figure 2A). We found significant main effects for nicotine dose [$F_{(2, 21)} = 20.35$, $p < 0.0001$], sex [$F_{(1, 21)} = 30.23$, $p < 0.0001$], chronic pain [$F_{(1, 21)} = 39.72$, $p < 0.0001$], and global interactions between these three [$F_{(2, 21)} = 8.470$, $p = 0.0020$]. Additionally, there were significant interactions between nicotine dose and chronic pain state [$F_{(2, 21)} = 5.166$, $p = 0.0150$], between nicotine dose and sex [$F_{(2, 21)} = 19.42$, $p < 0.0001$], and between chronic pain and sex [$F_{(1, 21)} = 14.93$, $p = 0.0009$].

Additional three-way ANOVA analyzed interactions between chronic pain state, nicotine dose, and sex effects (Supplementary Figure 2B). We found significant main effects for nicotine dose [$F_{(2, 21)} = 13.25$, $p = 0.0002$] and chronic pain [$F_{(1, 21)} = 34.95$, $p < 0.0001$], but no main effects of sex [$F_{(1, 21)} = 1.964$, $p = 0.1757$]. There was also no global interactions between these three [$F_{(2, 21)} = 0.9058$, $p = 0.4194$]. However, there were significant interactions between nicotine dose and chronic pain state [$F_{(2, 21)} = 4.230$, $p = 0.0286$] and between chronic pain and sex [$F_{(1, 21)} = 34.33$, $p < 0.0001$], but no interactions between nicotine dose and sex [$F_{(2, 21)} = 1.085$, $p = 0.3562$].

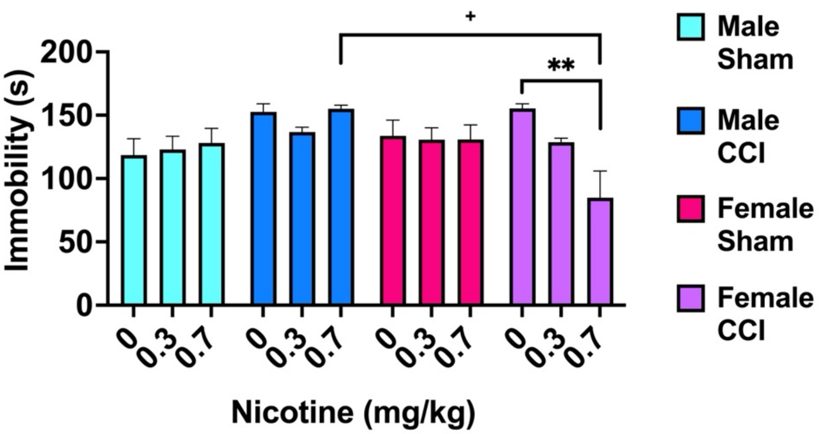


**Supplementary Figure 3: Sex differences in forced swim test immobility during nicotine withdrawal: three-way ANOVA.** A three-way ANOVA analyzed interactions between chronic pain state, nicotine dose, and sex effects (Supplementary Figure 3). No significant main effects were found for chronic pain [$F_{(1, 21)} = 2.943$, $p = 0.1009$], sex [$F_{(1, 21)} = 2.035$, $p = 0.1684$], and nicotine dose [$F_{(2, 21)} = 2.243$, $p = 0.1309$]. There was also no global interactions between these three [$F_{(2, 21)} = 1.793$, $p = 0.1910$]. However, there were significant interactions between nicotine dose and chronic pain state [$F_{(2, 21)} = 5.300$, $p = 0.0137$], and between chronic pain and sex [$F_{(1, 21)} = 5.258$, $p = 0.0323$], and interactions between nicotine dose and sex [$F_{(2, 21)} = 4.967$, $p = 0.0171$].

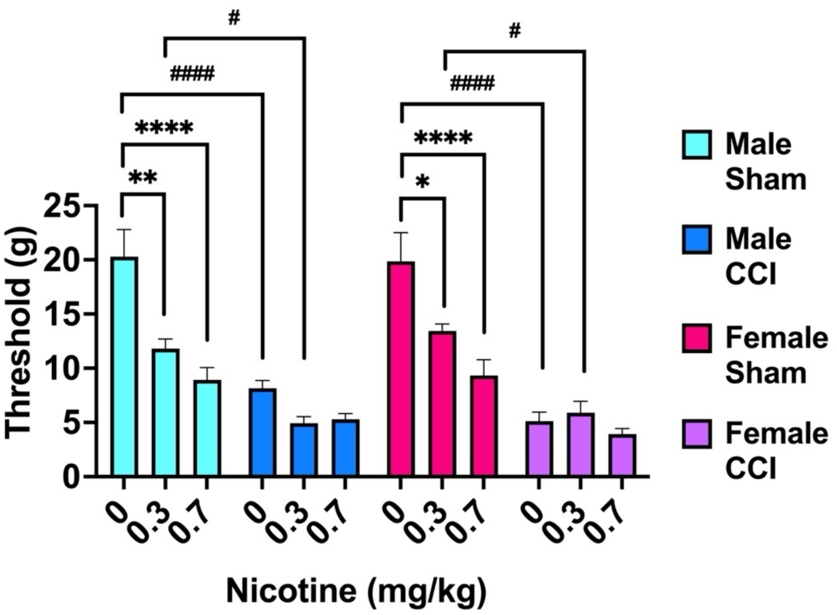


**Supplementary Figure 4: Sex differences in pain sensitivity on the withdrawal day: three-way ANOVA.** A three-way ANOVA evaluated the interactions of chronic pain state, nicotine dose, and sex on paw withdrawal thresholds. Significant main effects were observed for nicotine dose [$F_{(2, 21)}$ = 28.12, $p$ = 0.0001] and surgery [$F_{(1, 21)}$ = 207.4, $p$ < 0.0001], whereas sex had no significant effect [$F_{(1, 21)}$ = 0.1104, $p$ = 0.7430]. No global three-way interaction among surgery, dose, and sex was detected [$F_{(2, 21)}$ = 0.1119, $p$ = 0.8947]. A robust dose x surgery interaction indicated that the influence of nicotine dose differed between sham and CCI groups [$F_{(2, 21)}$ = 20.58, $p$ < 0.0001]. Neither dose x sex [$F_{(2, 21)}$ = 0.9737, $p$ = 0.3941] nor surgery x sex [$F_{(1, 21)}$ = 1.028, $p$ = 0.3221] interactions reached significance, indicating no evidence for sex-dependent modulation of mechanical sensitivity.

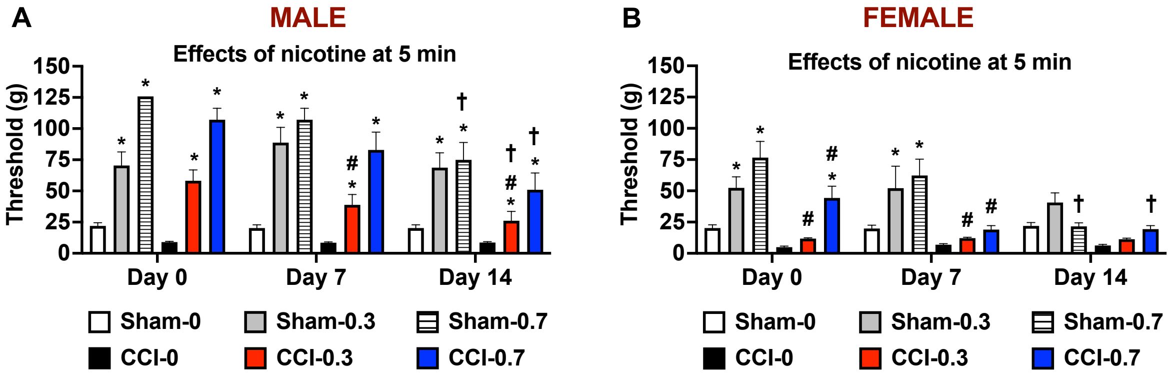


**Supplementary Figure 5. Sex differences in antinociceptive effects of nicotine in sham rats at 5 min: three-way ANOVA.** An additional three-way repeated-measures ANOVA evaluated the interactions between chronic pain state, nicotine dose, and sex, with time (day) included as a within-subject factor on paw withdrawal threshold at 5 minutes. Significant main effects were observed for chronic pain state [$F_{(1, 84)} = 72.13$, p < 0.0001], sex [$F_{(1, 84)} = 87.75$, $p < 0.0001$], nicotine dose [$F_{(2, 84)} = 109.93$, $p < 0.0001$], and time [$F_{(2, 168)} = 19.52$, $p < 0.0001$]. The interaction between chronic pain state and nicotine dose was significant [$F_{(2, 84)} = 5.02$, $p = 0.0087$], as was the nicotine dose x time interaction [$F_{(4, 168)} = 10.35$, $p < 0.0001$], indicating that the effects of nicotine dose varied over time. There was also a significant sex x nicotine dose interaction [$F_{(2, 84)} = 24.8478$, $p < 0.0001$]. No significant interactions were detected for chronic pain state x sex [$F_{(1, 84)} = 0.21$, $p = 0.651$], chronic pain state x sex x nicotine dose [$F_{(2, 84)} = 0.009$, $p = 0.991$], chronic pain state x time [$F_{(2, 168)} = 1.36$, $p = 0.260$], sex x time [$F_{(2, 168)} = 1.19$, $p = 0.306$], chronic pain state x sex x time [$F_{(2, 168)} = 2.01$, $p = 0.137$], chronic pain state x nicotine dose x time [$F_{(4, 168)} = 0.99$, $p = 0.417$], sex x nicotine dose x time [$F_{(4, 168)} = 0.20$, $p = 0.940$], or the four-way interaction among chronic pain state, sex, nicotine dose, and time [$F_{(4, 168)} = 1.18$, $p = 0.323$].
